# Enantiopurity-Dependent Peptide Coacervates and Asymmetric Organocatalysis

**DOI:** 10.1101/2025.08.13.670095

**Authors:** Alice Vetrano, Nico di Fonte, Olivier Monasson, Federico Perrella, Martina Porco, Ferdinando Mercuri, Gianluca Dell’Orletta, Francesco Petragnano, Alessio Carioscia, Davide Deodato, Damiano Calcagno, Giovanna Salvitti, Samantha Reale, Fabio Pesciaioli, Carino Ferrante, Davide Tedeschi, Giuseppe Grasso, Paola Benassi, Elisa Peroni, Armando Carlone, Isabella Daidone, Claudio Iacobucci

## Abstract

Membraneless compartmentalization via liquid–liquid phase separation (LLPS) has emerged as a powerful strategy to organize biochemical reactions. Recently, peptide-based coacervates demonstrated the potential to function as microreactors by enhancing reaction kinetics through increased local concentrations and altered microenvironments. Here, we introduce an O-methylated diphenylalanine-based tripeptide ^LLL^PFF-OCH_3_ containing an N-terminal proline, designed to undergo LLPS, and simultaneously function as an enantioselective organocatalyst. Comprehensive characterization via confocal microscopy, fluorescence recovery after photobleaching (FRAP), micro-Raman and attenuated total reflection infrared (ATR-IR) spectroscopy, diffusion-surface plasmon resonance (*D*-SPR), and molecular dynamics (MD) simulations revealed the formation of stable liquid droplets. In contrast, a racemic mixture of ^LLL^PFF-OCH_3_ and ^DDD^PFF-OCH_3_ failed to form liquid droplets and instead formed a solid precipitate, unveiling a critical role of enantiopurity in LLPS. Proof-of-concept catalytic studies proved enantioselective organocatalytic activity of the ^LLL^PFF-OCH_3_ liquid coacervates. Beyond catalysis these results may have broader implications in understanding prebiotic chemistry and neurodegeneration.

---

Biological systems rely on the spatial and temporal organization of chemical processes. While membrane-bound organelles serve this function in modern cells, many essential reactions occur in dynamic, membraneless compartments formed through liquid–liquid phase separation (LLPS).^[1]^ These biomolecular condensates serve to concentrate specific molecules and can modulate reaction kinetics and specificity.^[2]^ Bioinspired by these phenomenon, peptide-based coacervates have recently been introduced as synthetic analogues of biomolecular condensates.^[3]^ Composed of short sequences, such as *O*-methylated diphenylalanine derivatives,^[4]^ these coacervates can solubilize hydrophobic compounds and enzymes in water, acting as supramolecular microreactors.^[4,5]^ Previous studies have shown that encapsulating enzymes or catalysts within peptide droplets can accelerate chemical transformations due to local concentration effects and changes in rate constants.^[6]^

Proline-based peptides have been used in organocatalysis,^[7]^ with their self-assembly into gels and fibrils providing structural environments that modulate and support catalysis; indeed, we have recently shown that organocatalytic fibrils, self-assembled from minimal peptides, accelerate catalysis compared to their non-supramolecular organized counterparts.^[8,9]^

In this work, we introduce a tripeptide that both undergoes LLPS and is a competent enantioselective organocatalyst itself. By integrating L-proline at the N-terminus of an *O*-methylated diphenylalanine-based scaffold we developed the tripeptide ^LLL^PFF-OCH_3_, capable of forming chiral liquid droplets that function as asymmetric microreactors in aqueous environments. We characterized these systems at a molecular level using an integrative bioanalytical and computational approach, including confocal microscopy, FRAP, micro-Raman and ATR-IR spectroscopy, *D*-SPR, and MD simulations. Catalytic tests on a model aldol reaction revealed the enantioselective effect of the chiral coacervates. Strikingly, the racemic tripeptide failed to undergo LLPS, instead leading to immediate precipitation. This suggests that enantiopurity may be a prerequisite for spontaneous LLPS and compartmentalization in aqueous environments.

It is worth noting that the enantiopurity-dependent LLPS reported here involves the self-assembly of a single, short peptide. This is conceptually distinct from previous studies, such as those by Perry and co-workers, where LLPS was observed only when mixing oppositely charged polypeptides, and only if at least one was heterochiral, i.e., an optically inactive polymer with randomly distributed D- and L-monomers leading to a statistical mixture of diastereomers.^[10]^ In contrast, our system demonstrates that enantiopurity can be itself a prerequisite for LLPS of a single homochiral peptide.

## Results and discussion

Solubilizing ^LLL^PFF-OCH_3_ (Figure 1a) in water yielded a slightly acidic (pH ∼ 6), transparent solution (Figure 1b,c). Upon increasing the pH above 7.5, the solution turned turbid and white (Figure 1b,d). Microscopy experiments revealed that the turbidity corresponded to the LLPS of spherical coacervates formed by the concentrated tripeptide, with droplet diameters of approximately 5 my (Figure 1d,g). Droplet formation was fully reversible upon repeated pH cycling, indicating the dynamic nature of the phase separation process (Figure 1i).

**Figure 1.**
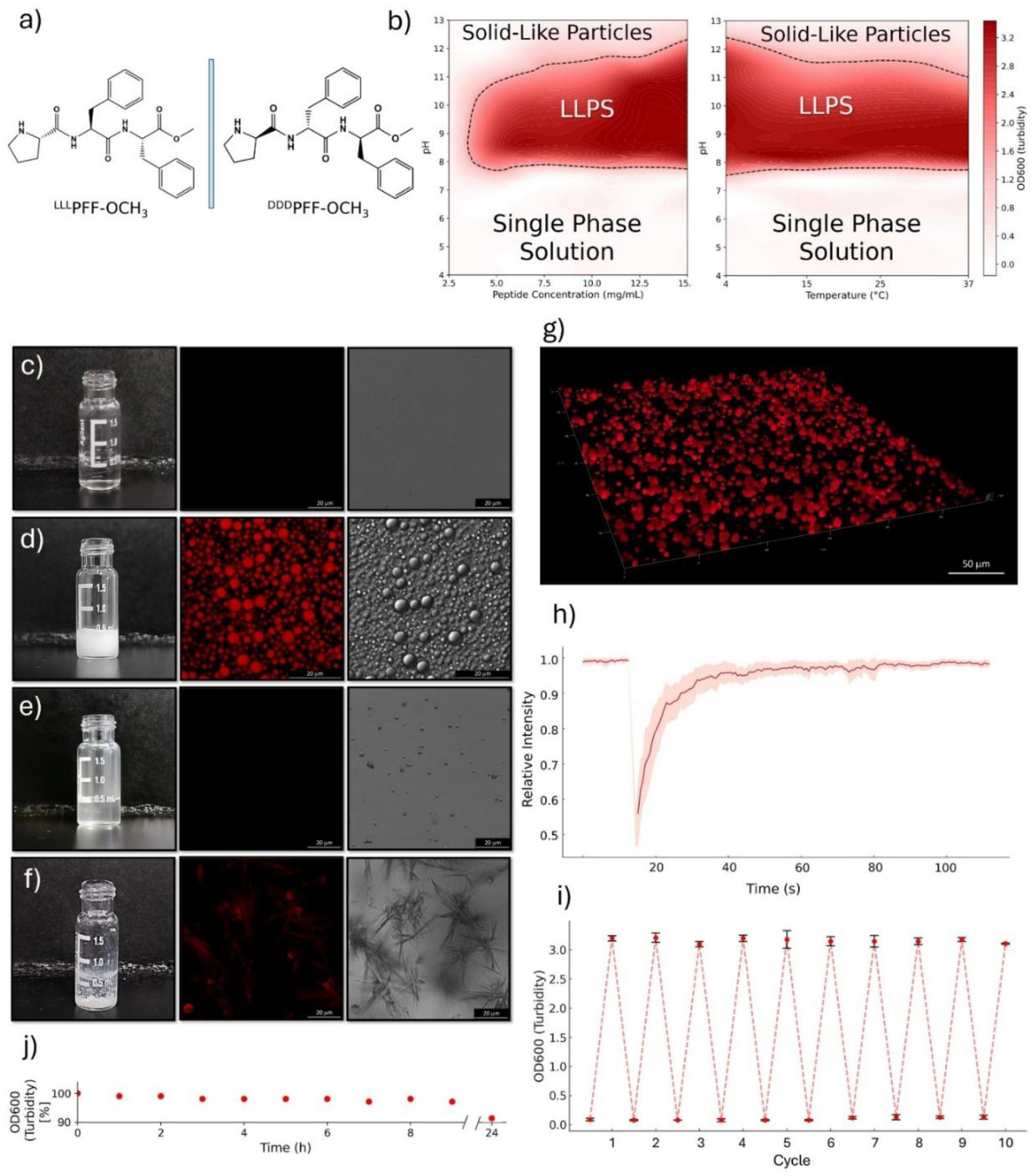
Phase behavior and morphology of PFF-OCH_3_ peptide. a) Chemical structure of ^LLL^PFF-OCH_3_ and ^DDD^PFF-OCH_3_. b) Phase diagrams depicting the formation of peptide droplets as a function of peptide concentration vs pH (left) and temperature vs pH (right), determined by turbidity (OD600). The LLPS region (dark red) is separated from the single phase solution and from the solid-like particles suspension by dashed boundaries. c–f) Vial appearance, autofluorescence confocal image, and micrograph (scale bar = 20 μm) of 10 mg mL^-1^ solutions of: c) ^LLL^PFF-OCH_3_ at pH 6; d) ^LLL^PFF-OCH_3_ at pH 8.5; e) ^LLL^PFF-OCH_3_ at pH 13; f) racemic (1:1) mixture of ^LLL^PFF-OCH_3_ and ^DDD^PFF-OCH_3_ at pH 8.5. g) 3D autofluorescence confocal reconstruction of ^LLL^PFF-OCH_3_ coacervates (10 mg mL^-1^, pH 8.5). h) FRAP plot of ^LLL^PFF-OCH_3_ coacervates (10 mg mL^-1^, pH 8.5) over time. The solid line and shaded area represent the mean and standard deviation over six replicates, respectively. i) Reversibility of LLPS over ten cycles of pH change between 6 and 8. Error bars represent the standard deviation of turbidity from three replicates. j) Stability of ^LLL^PFF-OCH_3_ coacervates (10 mg mL^-1^, pH 8.5, 20 °C) over time, monitored as the percentage of initial turbidity.

We systematically investigated the LLPS behavior of ^LLL^PFF-OCH_3_, evaluating its phase stability and responsiveness to changes in peptide concentration, pH, and temperature. To delineate the phase transition boundaries, we constructed concentration–pH and temperature–pH phase diagrams (Figure 1b). LLPS was observed at concentrations above 2.5 mg·mL^-1^ and within a pH range of ∼ 7.5 to 12, with lower temperatures further promoting phase separation. Above pH 12 the turbidity reduced and solid-like particles of ∼ 1 μm became visible (Figure 1e). These LLPS conditions are consistent with those reported for the LLPS of ^LL^FF-OCH_3_, suggesting that the presence of a C-terminal proline in the tripeptide has minimal influence on droplet formation.^[4]^

The LLPS mechanism proposed for ^LL^FF-OCH_3_ likely remains valid: attractive π–π interactions between the aromatic rings of the FF core drive assembly, while repulsive electrostatic interactions involving the partially positively charged N-termini help maintain solubility and define the pH responsiveness of the system. Experimental evidences supporting this hypothesis will be presented later in this paper.

Beyond the morphological characterization, we investigated the internal dynamics of the droplets using FRAP (Figure 1h). Complete fluorescence recovery within ∼ 60 seconds post-bleaching indicates high internal molecular mobility, consistent with the liquid-like nature of the coacervates.

The liquid droplets formed by ^LLL^PFF-OCH_3_ at pH 8.5 proved stable for days (Figure 1j) and under conditions typically disruptive for non-covalent peptide–peptide interactions, including 1 M NaCl, 2 M urea, and up to 40% acetonitrile, above which the turbid solution reverts to transparency. We applied D-SPR analysis;^[11,12]^ to characterize the diffusive behavior of 0.5 and 8 mg·mL^-1^ solutions of ^LLL^PFF-OCH_3_ at pH 6 and 8.5. At 0.5 mg·mL^-1^, below the coacervation threshold (Figure 1b), solutions remained transparent at both pH values, and a single diffusion coefficient of ∼ 3.5·10^−10^ m^2^·s^-1^ was observed (Figure S1), consistent with low-molecular-weight species. A comparable value was detected for the transparent 8 mg·mL^-1^ solution at pH 6. Notably, at 8 mg·mL^-1^ and pH 8.5, where coacervation occurs, an additional slower diffusing population emerged with a diffusion coefficient of ∼ 5.9·10^−11^ m^2^·s^-1^ (Figure S1), indicative of higher molecular weight species. These accounted for ∼ 20% of the total diffusion signal,^[13]^ suggesting that most peptides remain in minimal-size oligomers, consistent with the transient and dynamic interactions driving LLPS.^[14]^ We next investigated the effect of enantiopurity on the LLPS behavior of the tripeptide. Unlike proteins undergoing LLPS, which are inherently enantiopure, synthetic short peptides offer the opportunity to examine whether enantiomeric excess itself influences LLPS. To this end, we prepared a racemic mixture by combining equal volumes of ^LLL^PFF-OCH_3_ and ^DDD^PFF-OCH_3_, each at 10 mg·mL^-1^ in water at pH 6, resulting in a final solution containing 5 mg·mL^-1^ of each enantiomer (10 mg·mL^-1^ total peptide concentration). The resulting racemic solution remained transparent at pH 6, just like the two enantiopure solutions. Notably, this racemic mixture lies at the center of the LLPS region in the phase diagram for pH > 7.5, both in terms of total peptide concentration and individual enantiomer concentration. However, upon adding a precise amount of NaOH to adjust the solution to pH 7.5, the system did not undergo LLPS by turning turbid but instead underwent a liquid-to-solid transition, forming large, suspended white flocculent aggregates while remaining optically transparent in the bulk phase (Figure 1f). The racemic mixture obtained by combining equal volumes of ^LL^FF-OCH_3_ and ^DD^FF-OCH_3_, under the same concentration and condition, also did not underwent LLPS like the enantiopure ^LL^FF-OCH_3_^[4]^ but resulted in immediate precipitation. To further explore the role of enantiopurity, we prepared scalemic mixtures at 2:1 and 1:2 ratios of ^LLL^PFF-OCH_3_ and ^DDD^PFF-OCH_3_, keeping the total peptide concentration constant at 10 mg·mL^-1^. At pH 6, both solutions were transparent. Upon adjusting the pH to 7.5, the scalemic mixtures turned white and turbid, exhibiting suspended flocculent aggregates, similar to the racemic solution. Upon precipitation, the solid aggregates were separated by sedimentation, leaving a PFF-OCH_3_ excess in solution at 3.3 mg·mL^-1^, a concentration at which LLPS is expected (Figure 1b). Microscopy of the supernatants revealed the presence of spherical peptide droplets (Figure S2). This behavior is consistent with spontaneous enantiomeric self-disproportionation (SDE), a phenomenon in which scalemic mixtures separate into an enantioenriched solution and a racemic precipitate.^[14]^ We propose that this stereochemical fractionation represents an intrinsic property of the system, emerging independently of any catalytic function. As such, the observation of SDE in peptide-based LLPS is of fundamental interest in its own right and may bear implications for the origins of homochirality.^[14,15]^ The catalytic activity of the peptide droplets, which may further contribute to such an effects, will be discussed later in this manuscript.

Collectively, these results demonstrate that ^LLL^PFF-OCH_3_ undergoes robust and reversible LLPS above pH 7.5, forming liquids. Importantly, enantiopurity emerges as a critical factor: racemic and scalemic mixtures fail to undergo LLPS and instead exhibit spontaneous enantiomeric self-disproportionation, revealing a stereochemistry-based mechanism of phase separation.

Next, we investigated whether the conformational ensemble of ^LLL^PFF-OCH_3_ varies as a function of pH and enantiopurity. To this end, we adopted an integrative approach combining micro-Raman and IR spectroscopy, which provide complementary structural insights into both liquid and solid states. We recorded micro-Raman and IR spectra of ^LLL^PFF-OCH_3_ under three conditions: in solution before LLPS (pH 6), upon LLPS on single droplets (pH 8.5), and as a solid racemic mixture.

First, we compared the Raman spectrum of the transparent 10 mg·mL^-1^ solution of ^LLL^PFF-OCH_3_ at pH 6 with that recorded inside the droplets, upon coacervation of the same solution at pH 8.5 (Figure S3). The Raman mapping of a ^LLL^PFF-OCH_3_ droplet at pH 8.5 is reported in Figure 2a. The most prominent Raman feature was the sharp peak at 1003 cm^-1^, corresponding to the phenyl ring “breathing” mode. This signal is prominent even in diluted aqueous solutions and is commonly used to normalize Raman spectra of peptides and proteins due to its relative insensitivity to the chemical environment. We used the intensity ratio between this peak and the O–H stretching band of water at ∼ 3200 cm^-1^ to estimate the peptide concentration inside several droplets, which was 350±30 mg·mL^-1^ (758 mM) (see Supporting Information). This corresponds to a local concentration about 38 times higher than that of the initial solution at pH 6 (10 mg·mL^-1^, 22 mM), from which the droplets formed by increasing the pH to 8.5. This level of enrichment is consistent with the ∼ 75% water content reported for similar peptide-based coacervates.^[6]^ We then compared the six most intense Raman bands of the phenyl rings across the three aforementioned conditions. These signals, centered at 1606, 1586, 1207, 1031, 1003 and 622 cm^-1^, referred to as F1 to F6, are typically weakly sensitive to the molecular environment.^[16]^ As expected, their positions and intensities varied only slightly between the dilute pH 6 solution, the dense LLPS phase at pH 8.5, and the solid racemate. An exception was the 1207 cm^-1^ band (Figure 2b), whose vibrational mode involves the β-carbon of the phenylalanine side chain and is therefore more affected by the chemical environment. A progressive red shift across the three states suggests increased local packing and enhanced π–π interactions (Figure 2c). ^[16]^

**Figure 2.**
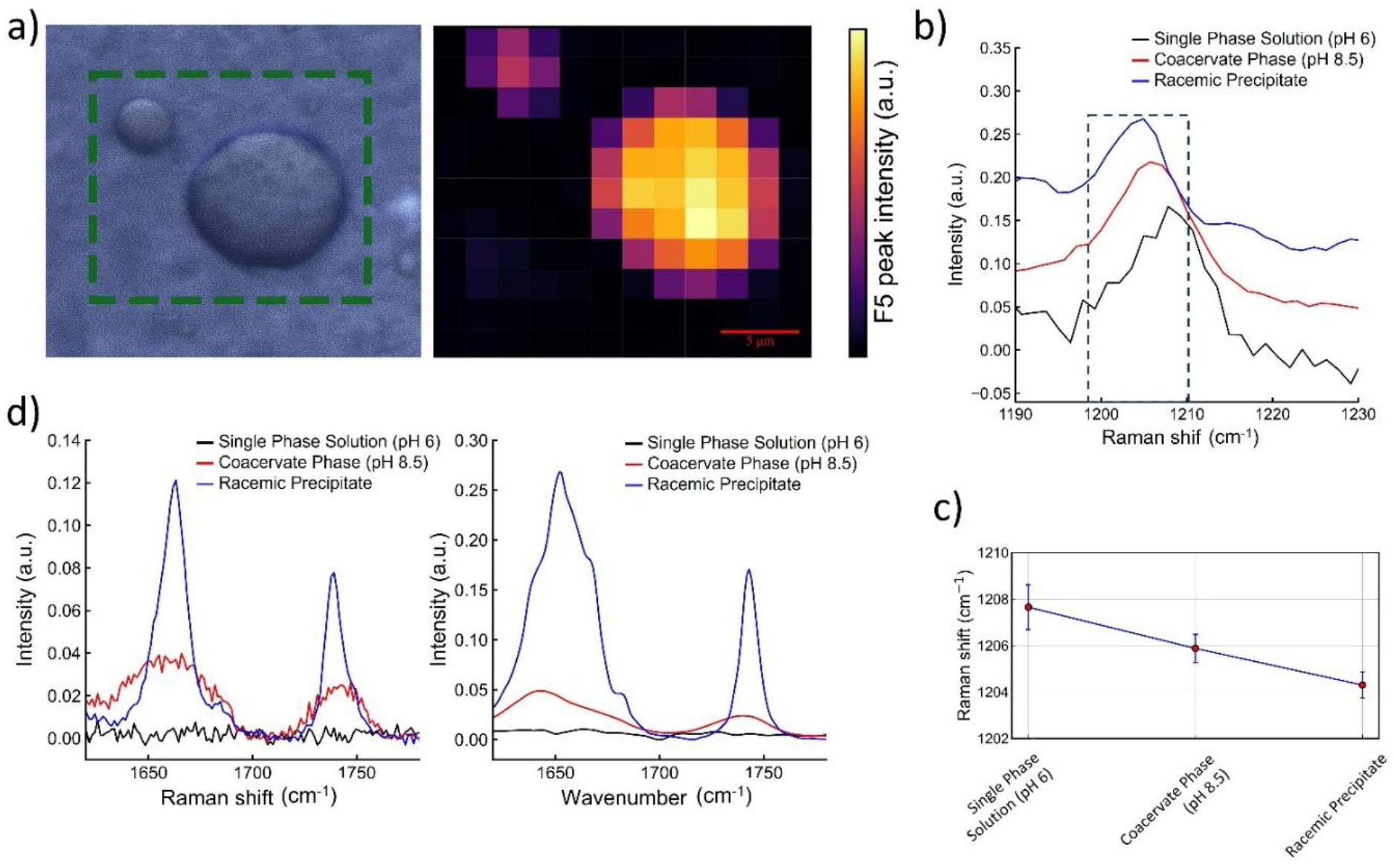
Spectroscopic characterization of ^LLL^PFF-OCH_3_ under different pH conditions. a) Optical micrograph (left) of an aqueous solution at pH 8.5, showing a representative coacervate droplet (green dashed box), and the corresponding Raman intensity map (right) of the boxed region for the F5 band (1003 cm^-1^).^[16]^ b) Raman spectra in H_2_O highlighting the F3 band for three sample states: single phase solution (pH 6, black), coacervate phase (pH 8.5, blue), and racemic precipitate (red). c) F3 peak positions (1207 cm^-1^) obtained by Gaussian fitting of the spectra in panel b); error bars indicate the 95% confidence interval of the fit. d) Comparison of Raman (left) and ATR-FTIR (right) spectra in D_2_O for the same three sample states, focusing on the C=O stretching vibrations of the amide I band and the methyl ester group (Raman: 1738 cm^-1^; IR: 1742 cm^-1^).

Next, we compared the peptide backbone conformation by analyzing the amide I region in both Raman and IR spectra (Figure 2d). To avoid overlap with the H2O bending band at 1640 cm^-1^, we used D2O-based solutions. Despite this, in the dilute pH 6 solution the amide I signal was indistinguishable from the noise. We therefore focused our comparison on the amide I region of the liquid droplets and the solid racemate. The IR spectrum of the enantiopure droplets displays a broad amide I band centered at 1645 cm^-1^ (FWHM ∼ 53 cm^-1^). Second-derivative analysis revealed three overlapping components centered at 1680, 1665, and 1641 cm^-1^, consistent with disordered structures, polyproline II (pPII)-like and β-strand/turn-like conformations.^[17–19]^ The corresponding Raman band displays a very broad amide I’ band centered at 1660 cm^-1^ (FWHM ∼ 51 cm^-1^) featuring at least three underlying components with maxima at 1686, 1667, 1651 cm^-1^, assigned to the same secondary structure elements.^[19–21]^ These features suggest that the peptide adopts a highly dynamic and disordered conformational ensemble within the droplets, consistent with their liquid nature and the transient character of interpeptide interactions. In contrast, the amide I band of the solid racemate is narrower (FWHM IR ∼ 37 cm^-1^, Raman ∼ 20 cm^-1^), with reduced contribution from disordered structures, in favor of β-strand signature. This shift reflects increased conformational order and rigidity in the solid racemate. The second peak present in the Raman (1738 cm^-1^) and IR (1742 cm^-1^) spectra is associated with the C=O stretching mode of the methyl ester. Also, in this case a narrowing of the FWHM is observed when moving from LLPS to the solid racemate.

Collectively, the spectroscopic data show that LLPS promotes a dynamic conformational ensemble enriched in disordered, pPII- and β-like structures, while the solid racemate adopts a more rigid and elongated conformations. This highlights how pH and enantiopurity modulate both phase behavior and peptide secondary structure.

To gain further molecular level understanding of the conformational ensemble and supramolecular organization of ^LLL^PFF-OCH_3_ coacervates we performed all-atom molecular dynamics (MD) simulations on the microsecond timescale (see the Supporting Information) mimicking the concentration (758 mM) inside the droplets experimentally estimated from micro-Raman data. This corresponds to a water-to-peptide ratio of approximately 75, specifically 538 peptide molecules and 39000 water molecules were used in the simulation box. To mimic the coacervate phase, the peptides were modelled as 93% cationic and 7% neutral, reflecting an estimated pH of ∼ 9.5, based on an assumed peptide pK_a_ of 10.6 typical of secondary amines. This mildly alkaline pH lies just above the experimentally tested range, ensuring a statistically relevant fraction of deprotonated peptides while remaining compatible with the LLPS phase diagram (Figure 1b). To maintain charge neutrality, an appropriate number of Cl^-^ counterions was added. To dissect the role of charge in supramolecular organization, two additional simulations were performed to represent the limiting cases of peptide protonation: one with fully protonated peptides, mimicking conditions well below the LLPS onset (Figure 1b), and another with fully deprotonated peptides, representing conditions beyond the LLPS stability range (Figure 1b,e). The fully protonated system reflects the behaviour of acidic, transparent peptide solutions, albeit at a higher concentration than in the corresponding experiment (10 mg·mL^-1^, 22 mM), while the fully deprotonated system captures the tendency of peptides to aggregate into insoluble, solid-like particles under strongly basic conditions. In all systems, the peptides were initially randomly placed in the simulation box, ensuring full hydration of the peptides.

Visual inspection of the MD trajectories reveals increased peptide aggregation with decreasing peptide charge, indicating that reduced electrostatic repulsion allows peptides to approach more closely and interact more extensively (Figure 3a–c). Cluster analysis performed using DBSCAN^[22–24]^ (Table S1) quantitatively shows that as the fraction of neutral peptides increases, the system transitions from many small, dispersed clusters to fewer, larger aggregates, ranging from 41 clusters on average with a mean size of 11 peptides in the fully cationic system to 13 clusters on average with a mean size of 47 peptides in the neutral system (Figure 3d-f, Figure S4-S6).

**Figure 3.**
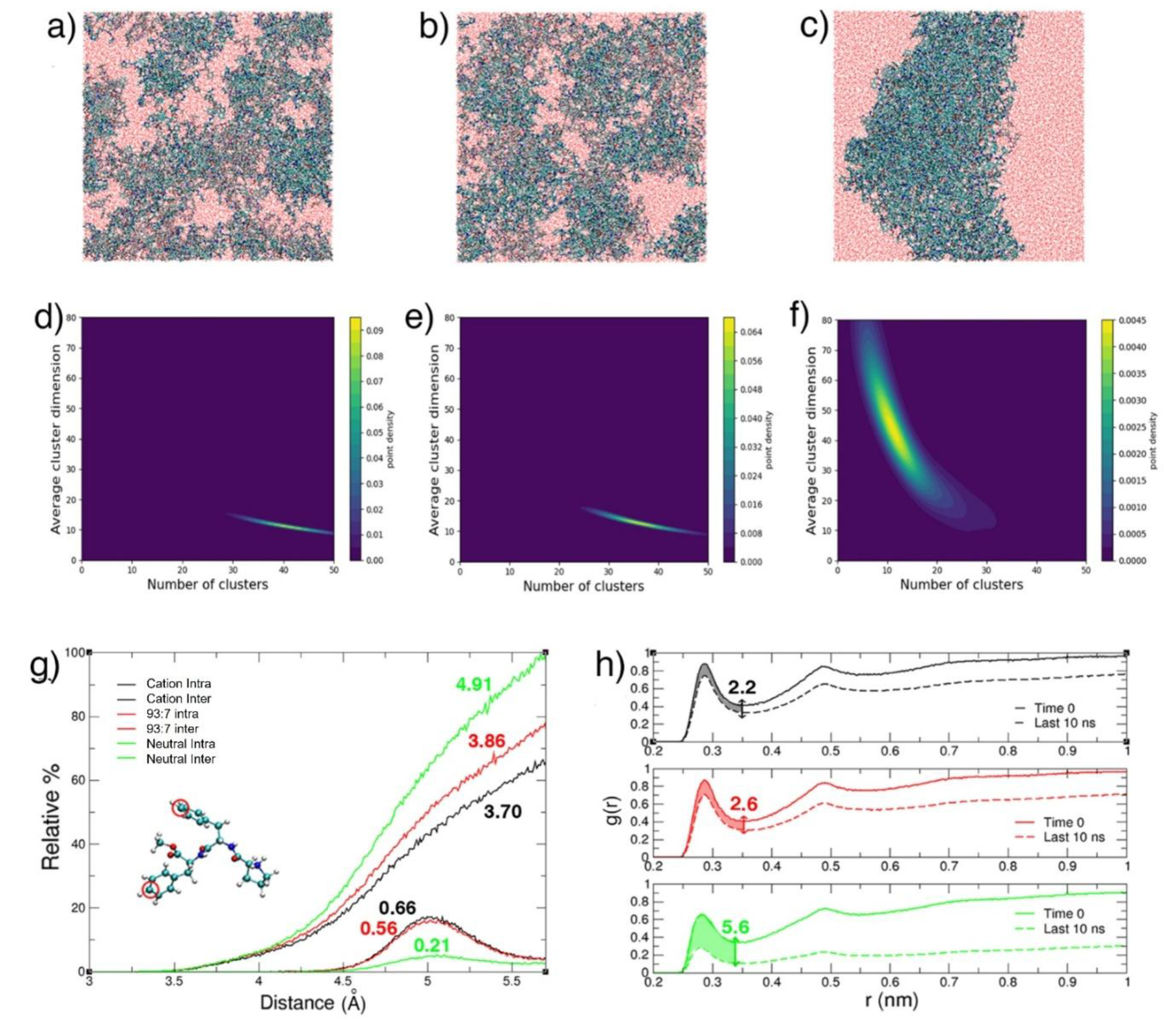
MD simulation results. Panels a–c: Representative snapshots from MD simulations illustrating peptide aggregation in the protonated, 93:7, and deprotonated systems, respectively. Panel d-f: Contour plots showing the correlation between the number of clusters and the average cluster size in the cationic, 93:7, and neutral systems, respectively. Panel g: Distribution of intra- and inter-molecular distances between the para carbon atoms (Cε) of phenylalanine side-chain phenyl rings, truncated at the first minimum of RDF, highlighting aromatic stacking interactions in the cationic (black), 93:7 (red) and neutral (green) systems. The values highlighted in the figure represent the average number of intramolecular and intermolecular interactions per molecule within MD simulation results. Radial distribution functions (RDFs) between peptide heteroatoms and water oxygen atoms are shown for the first nanosecond and the last ten nanoseconds of the simulation for each system (black lines represent the cationic system, red lines represent the 93:7 system, and green lines represent the neutral system). These RDFs are used to compare the decrease in hydration in the three systems. The difference in the coordination number of water molecules, integrated up to the first minimum, is indicated above the double arrow.

Analysis of distances between phenylalanine side chains (Figure 3g) reveals a clear increase in π–π stacking interactions between peptides as the concentration of neutral peptides rises. Quantitative assessment shows significantly more aromatic contacts within a 5.7 Å cutoff in systems containing neutral peptides, underscoring the critical role of aromatic interactions as the dominant driving force behind LLPS and aggregation of ^LLL^PFF-OCH_3_. Concurrently, intramolecular π–π stacking decreases. Interestingly, although the degree of aggregation is relatively high in all systems, the number of inter-peptide hydrogen bonds remains consistently low, amounting to 0.13, 0.15, and 0.47 per peptide in the fully cationic, 93:7 cationic:neutral, and fully neutral systems, respectively, (Figure S7) suggesting that aggregation is not primarily driven by hydrogen bonding.

Analysis of the φ/ψ dihedral angles (Figure S8) reveals that the backbone conformations cluster into three main basins: a broad basin centered at around –67°/140°, characteristic of polyproline II (pPII) conformations; a second basin centered at around –139°/150°, indicative of β-sheet–like conformations; and a third basin centered at around –66°/-43°, corresponding to turn-like conformations. Within the coacervate phase, the most populated basin is the pPII one, followed by the turn-like basin, with a minor contribution from the β-sheet–like basin. In the fully neutral system, the population of the β-sheet-like basin increases at the expense of the pPII and turn-like basins. These results indicate that the conformational repertoire within the coacervate phase is heterogeneous and largely disordered, lacking well-defined secondary structural elements, consistent with the observations from IR and Raman spectroscopy.

We finally examine the degree of peptide hydration in the coacervates and compare it to the fully cationic and fully neutral systems. Analysis of the radial distribution functions (RDFs) between peptide heteroatoms and water oxygens (Figure 3h) shows that hydration levels in the coacervates remain high, as evidenced by the pronounced first peak in the RDF, which is similar to that observed in the fully cationic system (Figure S9). The coordination number within the first hydration shell (up to the first minimum) is in fact 9.6 and 10.2 for the coacervate and fully cationic system, respectively. In contrast, comparison with the fully neutral system reveals a significant expulsion of water molecules from the peptide environment as neutrality increases, reflected by the overall decrease in the RDF intensity across the entire distance range. Specifically, the decrease in water coordination number from the fully dispersed state (i.e., at the start of the MD trajectories) to the equilibrated phase of the simulation becomes more pronounced when moving from the fully cationic system (in which the decrease is 2.2) to the fully neutral one (in which the decrease is 5.6).

These findings collectively support a model in which the reduced electrostatic repulsion resulting from an increased fraction of neutral peptides relative to cationic ones facilitates aromatic-driven aggregation, leading to fewer but larger peptide clusters. Altogether, these results highlight the critical interplay between peptide charge state, conformation, and supramolecular interactions that underpins the formation and stability of peptide-based coacervates.

Encouraged by the robust LLPS behavior of ^LLL^PFF-OCH_3_ and the ability of their droplets to solubilize organic molecules, such as rhodamine B during our FRAP experiments, we investigated whether these peptide droplets could function as confined chiral microreactors for enantioselective organocatalysis. In fact, rhodamine B exhibited a partition coefficient of 144:1 between ^LLL^PFF-OCH_3_ coacervate and the dilute phases (Figure S10), consistent with previous reports for ^LL^FF-OCH_3_ droplets.^[6]^ The prominent role that peptides have in asymmetric catalysis is majorly due to their unique combination of modularity and tunability. Synthetic peptides, particularly those of low molecular weight, can be easily prepared and customized by altering amino acid sequences, enabling fine-tuning of their structural and catalytic features.^[22]^ Their ability to adopt well-defined secondary structures allows for the precise spatial arrangement of functional groups, thereby promoting highly selective substrate activation^[23]^ and control over point, axial, and conformational chirality.^[24]^ Creating a chiral lipophilic microenvironment is indeed recognized as a good strategy to achieve (enantio)selective transformations.^[9]^ As a proof of concept, we tested the aldol reaction between cyclohexanone and benzaldehyde (Figure 4), a transformation known to be enantioselectively catalyzed by proline,^[25–27]^ and proline-containing peptides, with catalytic efficiency often depending on their supramolecular organization.^[9,28]^

**Figure 4.**
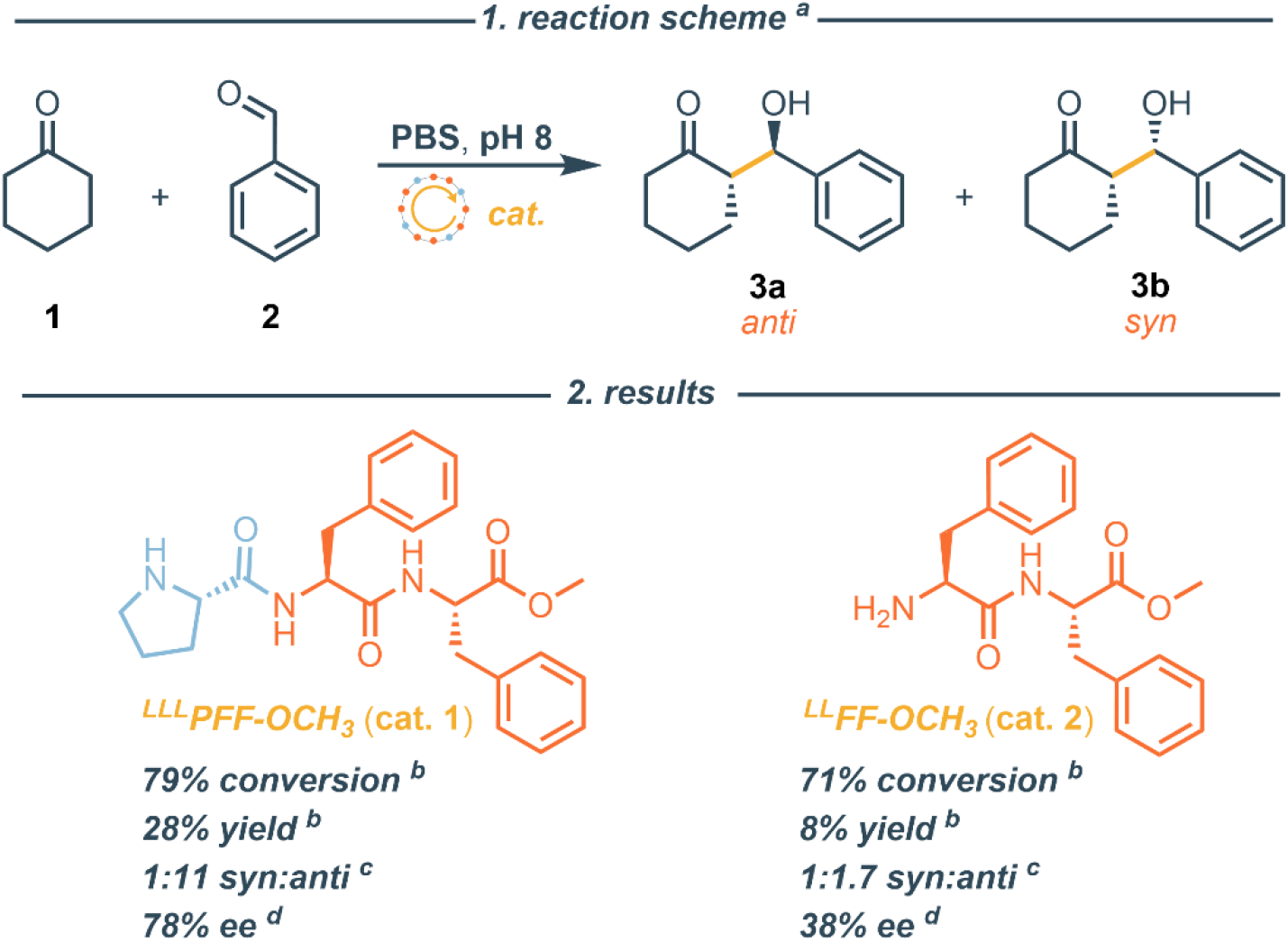
Peptide catalysed aldol reaction. 1. reaction scheme; 2. results of the catalysed reaction. [a] All reactions were performed with **2** (11.6 mg, 0.1 mmol, 1 equiv), **1** (21.4 mg, 0.2 mmol, 2 equiv), and **cat**. (0.02 mmol, 0.2 equiv) into a PBS aqueous solution (1 mL, 0.1 M) at pH 8.5, for 20 h at 4 °C. [b] Determined by ^1^H NMR using triphenylmethane as internal standard. [c] Diasteromeric ratio of the syn:anti β-hydroxy ketone **3a** and **3b** product determined by ^1^H NMR analysis of the crude mixture. [d] Enantiomeric excess of the major anti diasteroisomer **3a** determined by chiral HPLC analysis.

We compared the catalytic properties of enantiopure ^LLL^PFF-OCH_3_ droplets to the droplets formed from the ^LL^FF-OCH_3_ dipeptide. Reactions were conducted using 20 mol% of catalyst at pH 8.5 and 4 °C for 20 hours to ensure droplet density and stability. The ^LL^FF-OCH_3_ droplets proved competent for catalyzing the aldol reaction, in agreement with previous reports on similar ^LL^FF-OCH_3_-based coacervates.^[6]^ Interestingly, HPLC analysis of the β-hydroxy ketone product revealed that ^LL^FF-OCH_3_ droplets, despite lacking proline functionality, but thanks to a free amino group, can function as chiral microreactors, albeit with moderate enantioselectivity (ee= 38%). Notably, the ^LLL^PFF-OCH_3_ droplets outperformed the ^LL^FF-OCH_3_ ones in both diastereo- and enantioselectivity (Figure 4). We attribute this enhancement to the presence of the N-terminal proline, which promotes the formation of more sterically hindered enamine intermediates, offering improved stereochemical control^[29]^

In short, these results highlight that ^LLL^PFF-OCH_3_ coacervates encapsulate organic molecules enabling asymmetric organocatalysis where N-terminal proline enhance the stereochemical control.

In conclusion, our study demonstrates that the tripeptide ^LLL^PFF-OCH_3_ forms liquid droplets through enantiopurity-dependent LLPS, revealing spontaneous stereochemical self-disproportionation. Spectroscopic analyses and molecular simulations unveiled that droplet formation relates to a dynamic and heterogeneous conformational ensemble and to hydrophobic-driven interactions, modulated by the pH. Remarkably, these peptide droplets function as chiral microreactors, for enantioselective aldol reaction. Our findings highlight a fundamental link between molecular chirality and the spontaneous occurrence of LLPS of peptides in aqueous environments. The enantiopurity dependence observed for ^LLL^PFF-OCH_3_ and ^LL^FF-OCH_3_ LLPS suggests that it may be an essential prerequisite for the formation of stable biomolecular coacervates. This, together with the enantioselective catalytic capabilities of these liquid coacervates, is particularly intriguing from a prebiotic chemistry perspective, as early compartmentalization and selective molecular enrichment, driven by chirality-dependent LLPS, could have played a role in the emergence of homochirality.^[3,30]^ Furthermore, the chirality-sensitive LLPS we describe may offer significant implications for understanding pathological processes in neurodegenerative diseases. Dysregulated LLPS and protein aggregation contribute to the pathology of disorders such as Alzheimer’s and Parkinson’s diseases.^[31,32]^ Aberrant chirality at the amino acid level has been increasingly recognized as a pathological hallmark in these conditions. Literature also indicates that altered chirality can modulate peptide and protein interactions.^[33,34]^ Our results, showing that enantiopurity affects peptide LLPS behavior, suggest that similar chirality-dependent mechanisms might underline protein misfolding and aggregation in neurodegenerative pathologies.

## Supporting Information

Additional figures, tables, and a detailed description of the materials and methods are available in the Supporting Information.

## Acknowledgements

Armando Carlone and Claudio Iacobucci acknowledge funding by the European Union - NextGenerationEU under the Italian Ministry of University and Research (MUR) National Innovation Ecosystem grant ECS00000041 - VITALITY - CUP E13C22001060006. Alessio Carioscia acknowledges Dipharma Francis for supporting his PhD fellowship. Martina Porco acknowledges Fresenius Kabi iPSUM for supporting her PhD fellowship. Claudio Iacobucci acknowledges financial support by the Italian Ministry of University and Research (MUR) (PRIN 2022 – Project 20225HNCZK) and the European Union Next Generation EU (PRIN 2022 PNRR - Project P20224WAME). Armando Carlone acknowledges financial support by the European Union Next Generation EU (PRIN 2022 PNRR - Project P2022YM7F2).

