## supplementary information for "Enantiopurity-Dependent Peptide Coacervates and Asymmetric Organocatalysis"

[e] Dr. F. Petragnano
Department of Biotechnological and Applied Clinical Sciences, University of L'Aquila, Via Vetoio, L'Aquila, 67100, Italy

[f] Prof. G. Grasso, Dr. Davide Deodato
Department of Chemical Sciences, University of Catania, Viale Andrea Doria 6, 95125, Catania, Italy

[g] D. Calcagno
IRCCS-Fondazione Bietti, Rome, Italy

[h] Dr C. Ferrante
CNR- SPIN, c/o Department of Physical and Chemical Sciences, University of L'Aquila, Via Vetoio, L'Aquila, 67100, Italy

### These authors equally contributed

**Table of contents**


| | |
|---|---|
| **1. Supporting Figures** | **3** |
| Figure S1 | 3 |
| Figure S2 | 3 |
| Figure S3 | 4 |
| Figure S4 | 4 |
| Figure S5 | 5 |
| Figure S6 | 5 |
| Figure S7 | 6 |
| Figure S8 | 6 |
| Figure S9 | 7 |
| Figure S10 | 7 |
| **2. Synthesis and Characterization of PFF-$OCH_3$** | **8** |
| 2.1 Materials | 8 |
| 2.2 Synthesis of FF-$OCH_3$ and PFF-$OCH_3$ | 8 |
| 2.3 NMR Characterization | 12 |
| **3. Peptide Coacervates: Preparation and Functional Properties** | **16** |
| 3.1 Coacervate Preparation | 16 |
| 3.2 Turbidity Measurements | 16 |
| 3.3 Guest Molecule Partitioning | 17 |
| 3.4 D-SPR Analysis | 17 |
| **4. Characterization of Coacervates** | **17** |
| 4.1 Fluorescence Confocal Microscopy | 17 |
| 4.2 FRAP Measurements | 18 |
| 4.3 Micro-Raman Spectroscopy | 18 |
| 4.4 IR Spectroscopy | 19 |
| **5. Computational Analysis** | **20** |
| 5.1 Molecular Dynamic Details | 20 |
| 5.2 Computational Analysis (Table S1) | 20 |
| **6. Aldol Reactions in Coacervates** | **22** |
| 6.1 General Methods | 22 |
| 6.2 Experimental procedures | 23 |
| **7. Supplementary References** | **27** |

#### 1. Supporting Figures

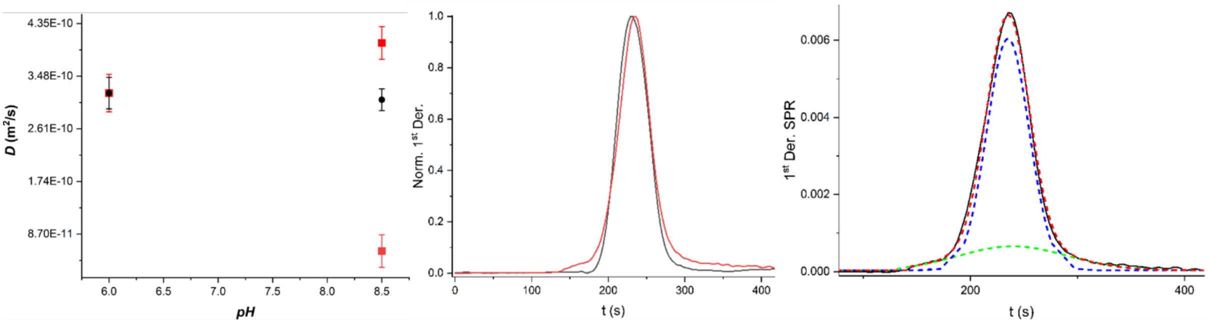


**Figure S1.** D-SPR analysis of $^{LLL}$PFF-$OCH_3$ diffusion**.** Left panel: Diffusion coefficients (D) of 0.5 mg·mL$^{-1}$ (black circles) and 8 mg·mL$^{-1}$ (red squares) peptide solutions at pH 6 and 8.5. Error bars indicate standard deviation from three independent experiments. At 8 mg·mL$^{-1}$ and pH 8.5, the D-SPR signal was deconvoluted into two peaks, corresponding to species with distinct molecular weights. Central panel: First derivative D-SPR signals for 8 mg·mL$^{-1}$ solutions at pH 6 (black line) and pH 8.5 (red line). Right panel: Deconvolution of the first derivative D-SPR signal at 8 mg·mL$^{-1}$ and pH 8.5 (black line). The red dotted line represents the cumulative fit; blue and green dotted lines indicate contributions from low and high molecular weight species, respectively.

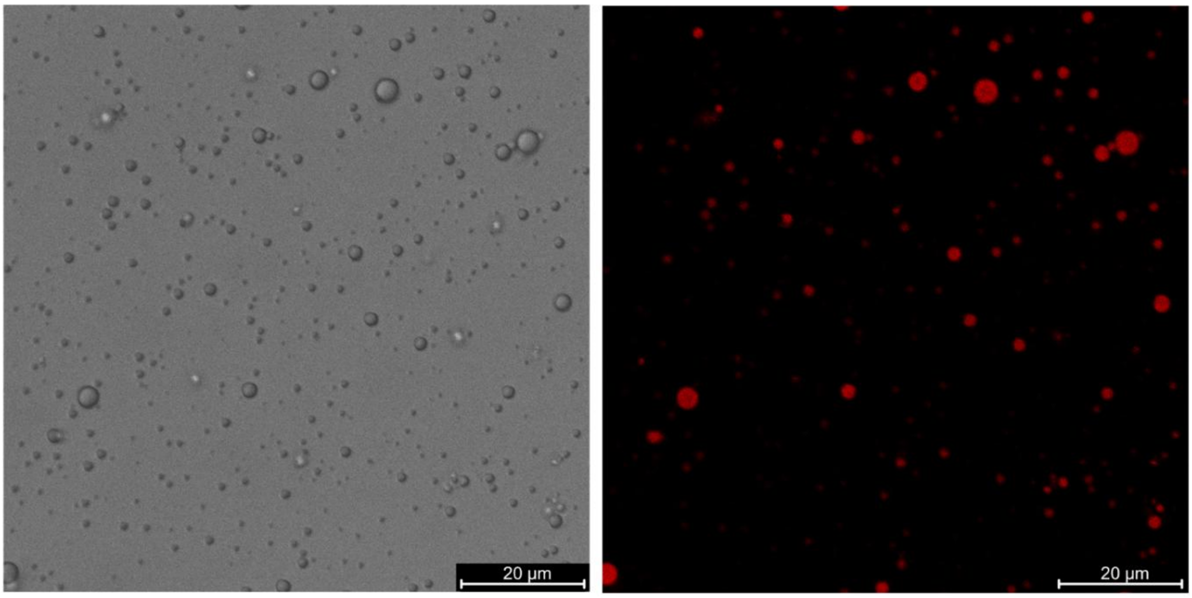


**Figure S2.** Brightfield micrograph and autofluorescence confocal image (scale bar = 20 μm) of the scalemic mixtures at 2:1 of $^{LLL}$PFF-$OCH_3$ and $^{DDD}$PFF-$OCH_3$ upon precipitation of the racemic component. The solid aggregates (Figure 1f) were removed by sedimentation. Microspopy revealed the presence of liquid droplets in the supernatant formed by the remaining $^{LLL}$PFF-$OCH_3$ enantiomer (3.3 mg·mL$^{-1}$).

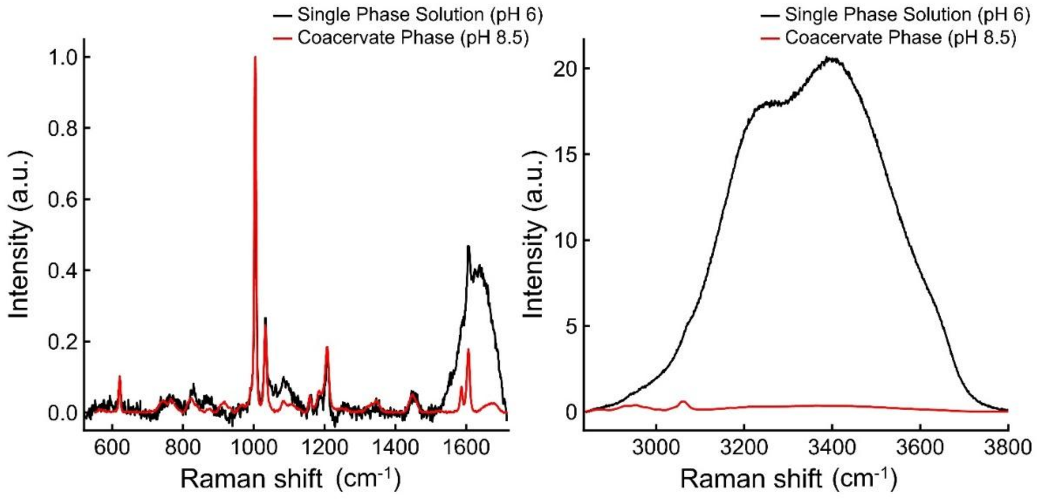


**Figure S3.** Normalized Raman spectra acquired inside a $^{LLL}$PFF-$OCH_3$ droplet at pH 8.5 (red) and for the transparent 10 mg·mL$^{-1}$ solution of $^{LLL}$PFF-$OCH_3$ at pH 6 (black). The left panel shows the fingerprint region (600–1800 cm$^{-1}$), highlighting enhanced signals inside the coacervates, including a sharp phenylalanine ring breathing band at ~ 1003 cm$^{-1}$ (F5). The right panel displays the high-wavenumber region (2800–3800 cm$^{-1}$), where a marked decrease in the broad OH stretching band is observed inside the coacervates, indicating reduced water content compared to the 10 mg·mL$^{-1}$ solution of $^{LLL}$PFF-$OCH_3$ at pH 6. The spectra are normalized to the band at ~ 1003 cm$^{-1}$.

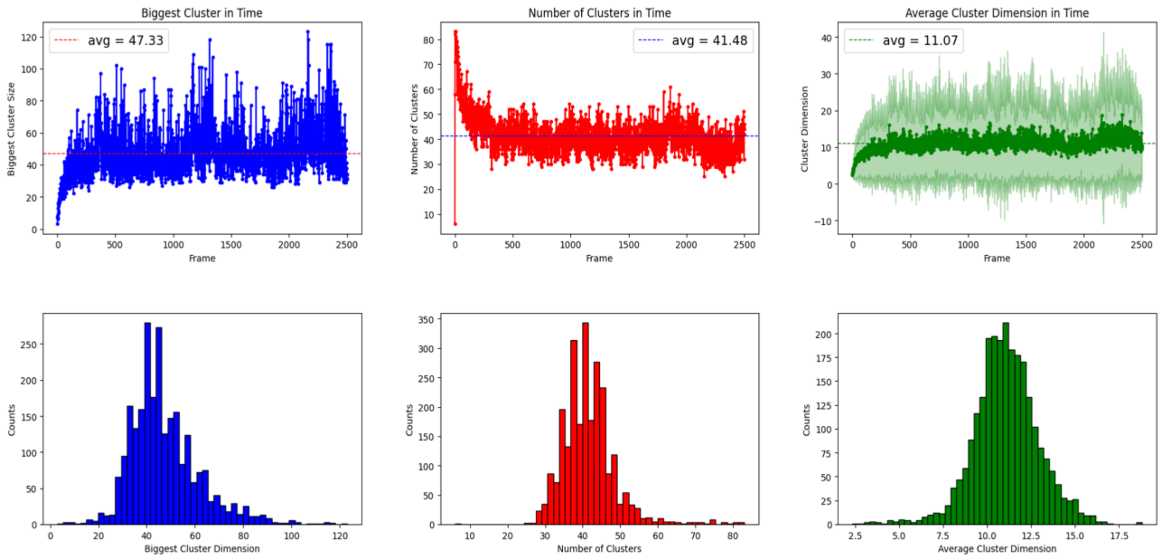


**Figure S4.** Cluster analysis of the fully cationic system. The plots show the size of the largest cluster over time, the total number of clusters over time, the average cluster size over time, and the distribution of cluster sizes.

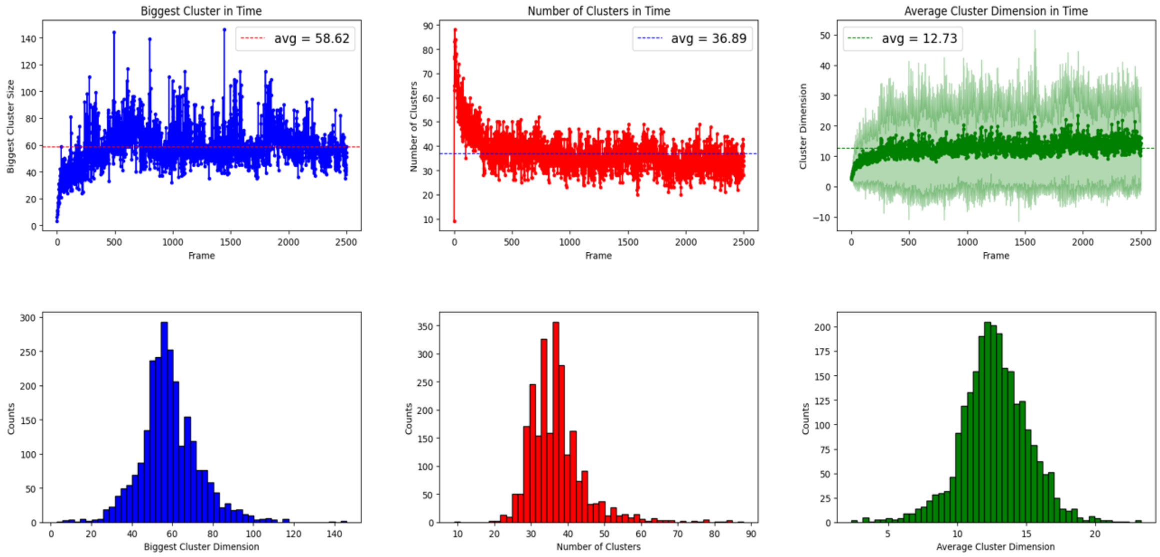


**Figure S5.** Cluster analysis of the system at pH ~ 9.5 (93:7 protonated:neutral ratio). The plots show the size of the largest cluster over time, the total number of clusters over time, the average cluster size over time, and the distribution of cluster sizes.

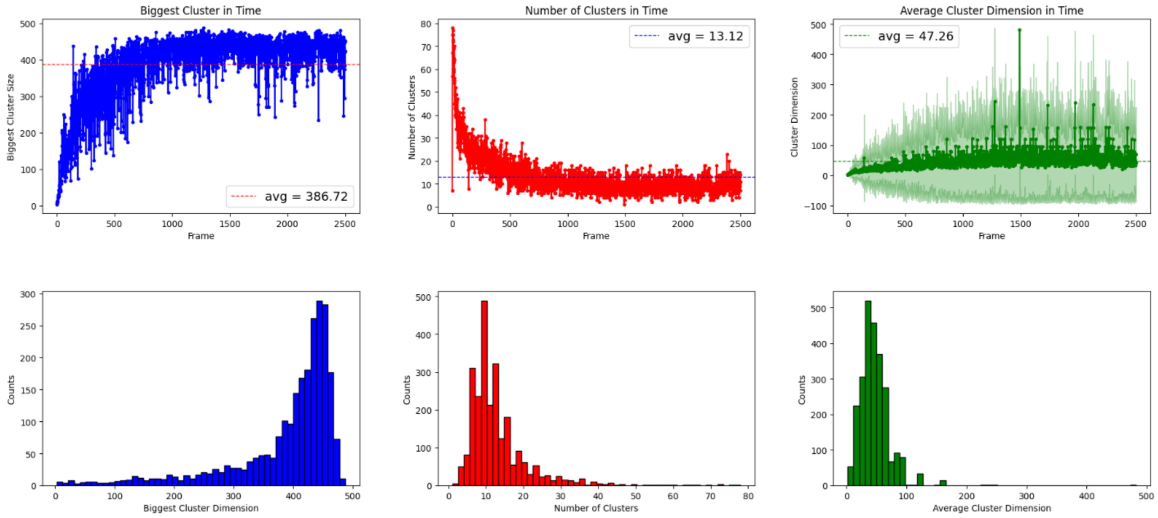


**Figure S6.** Cluster analysis of the fully deprotonated system. The plots show the size of the largest cluster over time, the total number of clusters over time, the average cluster size over time, and the distribution of cluster sizes.

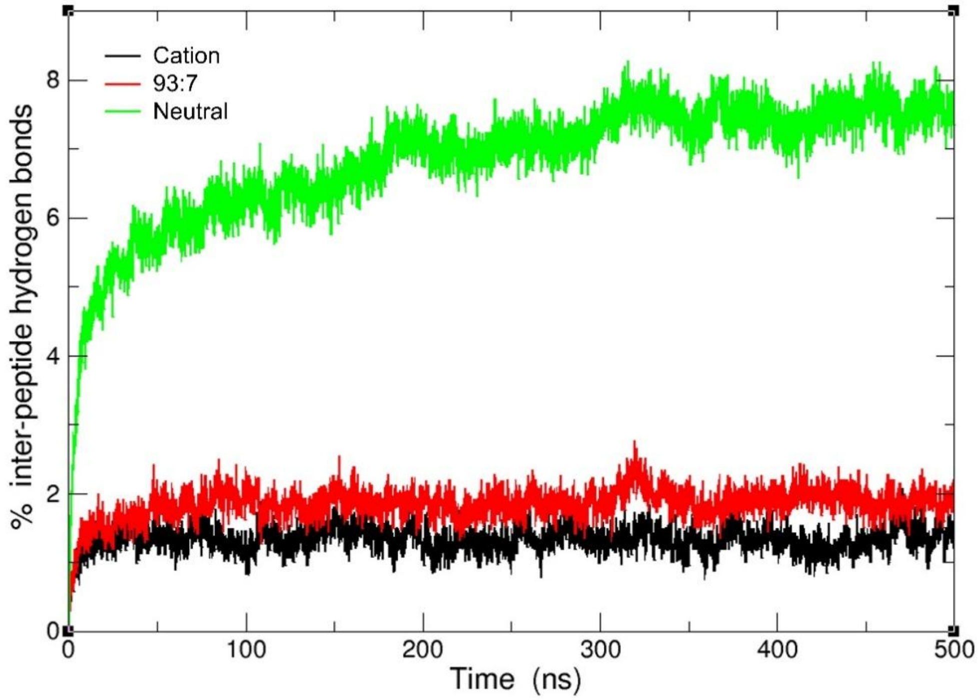


**Figure S7.** Time evolution of inter-peptide hydrogen bonding under different pH conditions. The percentage of inter-peptide hydrogen bonds is shown over time for the fully cationic system (black line), the partially deprotonated system at pH ~ 9.5 (93:7 protonated:neutral ratio, red line), and the fully neutral system (green line). A value of 100% corresponds to one hydrogen bond per potential donor or acceptor heteroatom.

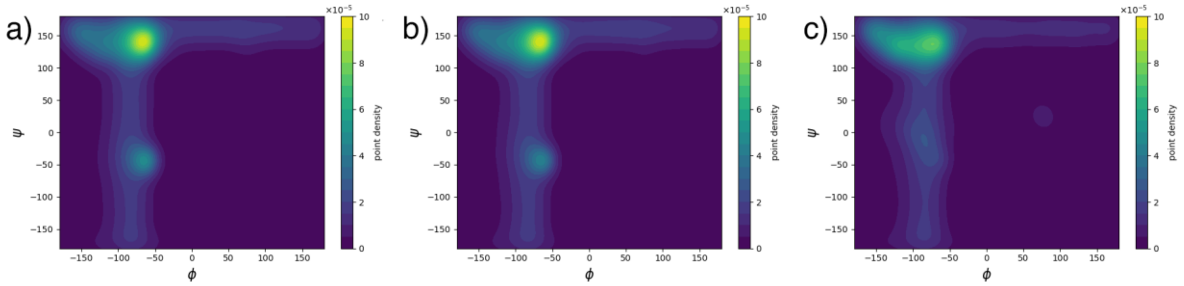


**Figure S8**. Ramachandran plots of $^{LLL}$PFF-$OCH_3$ under different pH conditions. Backbone dihedral angle distributions obtained from molecular dynamics simulations at: a) fully cationic state, b) pH ~ 9.5 (93:7 ratio of protonated to deprotonated peptides), and c) fully neutral state. The plots highlight how changes in protonation state modulate the conformational landscape of the peptide backbone.

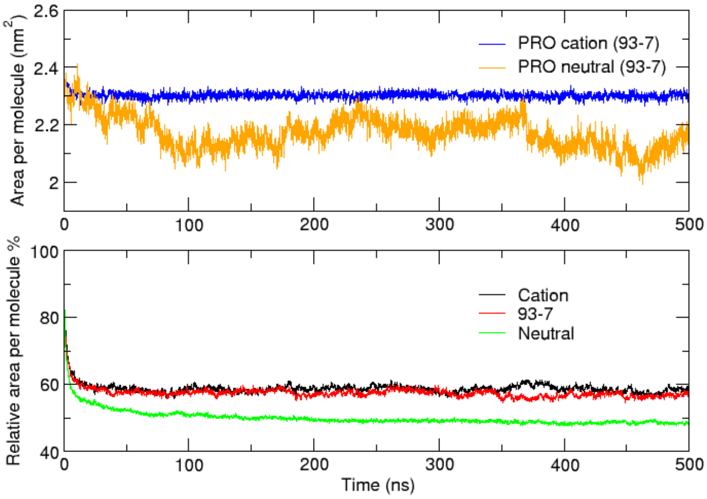


**Figure S9**. Time evolution of the relative solvent-accessible surface area (SASA) per molecule. Molecular dynamics simulations at pH ~ 9.5 (93:7 ratio of protonated to neutral $^{LLL}$PFF-$OCH_3$) show the evolution of SASA over time. The upper panel distinguishes the contributions from protonated and neutral prolines, highlighting differences in solvent exposure between the two species.

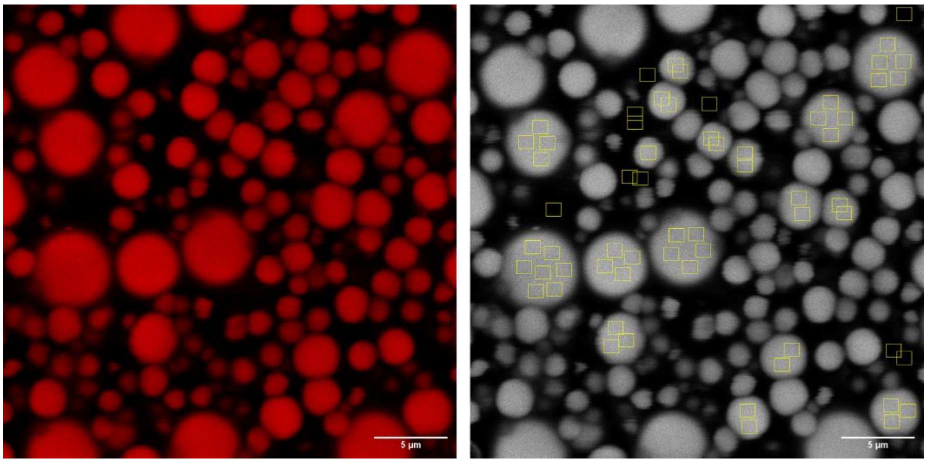

**Figure S10.** Rhodamine B partitioning into $^{LLL}$PFF-$OCH_3$ liquid droplets. Confocal fluorescence microscopy image showing selective accumulation of Rhodamine B (red) within the liquid droplets. A grayscale version of the image (right) highlights regions of interest (ROIs) placed inside and outside the droplets for analysis. The partition coefficient was calculated as the ratio of mean fluorescence intensity (mean gray value) between droplet and background ROIs. Scale bar = 5 µm.

**Synthesis and Characterization of PFF**

**2.1 Materials**

Boc-L-phenylalanine (Boc-L-Phe-OH, Iris Biotech), L-Phenylalanin-methylester hydrochlorid (L-Phe-OMe, Iris Biotech), Boc-L-proline (Boc-L-Pro-OH, Iris Biotech), Boc-D-phenylalanine (Boc-D-Phe-OH, Iris Biotech), D-Phenylalanin-methylester hydrochlorid (D-Phe-OMe, Iris Biotech), Boc-D-proline (Boc-D-Pro-OH, Iris Biotech), 1-hydroxybenzotriazole (HOBt, 97%, Sigma), N,N-diisopropylethylamine (DIPEA, 99%, Merck), 4 M hydrogen chloride solution in dioxane (Merck), N-(3-Dimethylaminopropyl)-N′-ethylcarbodiimide hydrochloride (EDC·HCl, Merck), Rhodamine B (Merck), CDCl3 (99.9% D, Merck), DMSO-d6 (99.9% D, Merck), phosphate buffered saline (PBS, Sigma). All the other solvents, chemicals, and salts used were purchased from Merck and were used as received.

**2.2 Synthesis of FF-OCH$_3$ and PFF-OCH$_3$**

$^{LL}$FF-OCH$_3$ and $^{LLL}$PFF-OCH$_3$ were synthesized following well-established multi-step procedures, with slight modifications to previously reported methods.[1,2] The corresponding synthetic schemes are shown below. Analogous strategies were employed for the preparation of $^{DD}$FFOCH$_3$ and $^{DDD}$PFF-OCH$_3$, affording similar yields.

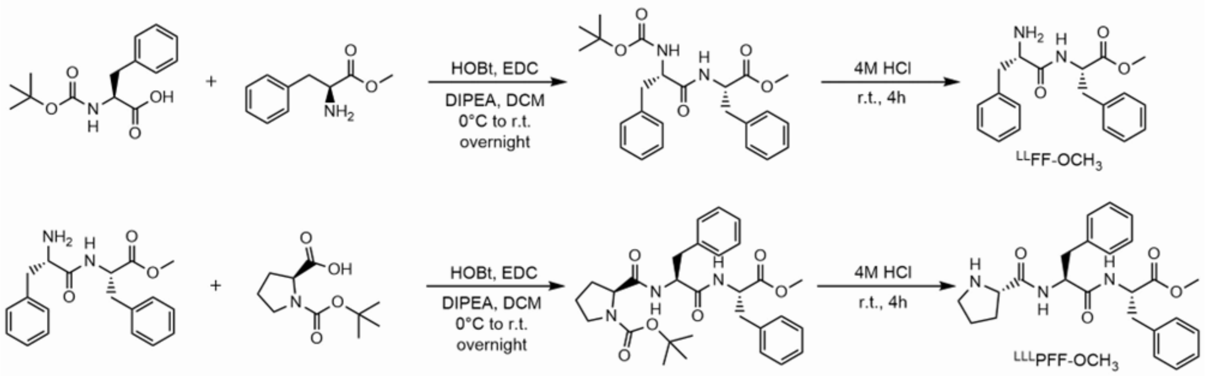


*N-Boc-L-phenylalanine-L-phenylalanine methyl ester:* N-Boc-L-phenylalanine (1.0 g, 4 mmol, 1.0 equiv.) was dissolved or suspended in dichloromethane (DCM) and cooled to 0 °C. To the resulting mixture were added 1-hydroxybenzotriazole monohydrate (HOBt, 4.4 mmol, 1.1 equiv.) and, after stirring for 5 min, N-(3-dimethylaminopropyl)-N′-ethylcarbodiimide hydrochloride (EDC·HCl, 4.4 mmol, 1.1 equiv.). The solution was stirred at 0 °C for 1 h. In parallel, L-phenylalanine methyl ester hydrochloride (0.93 g, 4.4 mmol, 1.1 equiv.) was dissolved in DCM and treated with N,N-diisopropylethylamine (DIPEA, 12 mmol, 3.0 equiv.) added dropwise at room temperature. After stirring for 1 h, this solution was added to the activated Boc-amino acid mixture, rinsing the flask with a small amount of DCM to ensure full transfer. The combined reaction mixture was stirred overnight at room temperature. Upon completion (monitored by TLC, petroleum ether/ethyl acetate 1:2), the reaction mixture was washed sequentially with 0.1 M HCl (3×), then with brine. The organic phase was dried over anhydrous $MgSO_4$, filtered, and concentrated under reduced pressure. The crude product was purified by flash chromatography on silica gel using a gradient elution of petroleum ether/ethyl acetate from 80:20 to 50:50. The pure product was obtained as a white foam (1.42 g, 3.48 mmol, yield: 87%) which was characterized by NMR. N-Boc-$^{LL}$FF-$OCH_3$ :$^1$H NMR (400 MHz, $CDCl_3$, 303 K) δ 7.53 – 7.10 (m, 10H), 7.00 (dd, J = 7.1, 2.3 Hz, 2H), 4.80 (d, J = 6.4 Hz, 1H), 4.36 (d, J = 6.2 Hz, 1H), 3.69 (s, 3H), 3.29 – 2.88 (m, 4H), 1.42 (s, 9H).

*L-phenylalanine-L-phenylalanine methyl ester*: The intermediate compound N-Boc-L-phenylalanine-L-phenylalanine methyl ester (1.42 g) was dissolved in 6 mL of 4 M hydrogen chloride in dioxane and stirred at room temperature for 3 hours to allow deprotection. The solvent was then removed under reduced pressure, affording an oily residue. Cold diethyl ether was added to the residue, resulting in the formation of a white precipitate. The supernatant was decanted, and the addition of diethyl ether followed by evaporation was repeated twice to ensure complete removal of residual reagents and solvents. The resulting white solid (1.1 g, yield: 99%) was collected and characterized by NMR and mass spectrometry. $^{LL}$FF-$OCH_3$ :$^1$H-NMR (400 MHz, $D_2O$, 303 K) δ 7.56 – 6.98 (m, 10H), 4.61 (dd, *J* = 8.2, 6.4 Hz, 1H), 4.09 (t, *J* = 7.1 Hz, 1H), 3.61 (s, 3H), 3.17 – 2.85 (m, 4H). HRMS (ESI-Q-Orbitrap): m/z calcd for $C_{19}H_{22}N_2O_3$ $[M+H]^+$ m/z = 327.1703, found m/z = 327.1698.

*N-Boc-L-proline-L-phenylalanine-L-phenylalanine methyl ester:* N-Boc-L-proline (0.65 g, 3.0 mmol, 1.0 equiv.) was dissolved or suspended in dry dichloromethane (DCM) and cooled to 0 °C. To the resulting mixture were added 1-hydroxybenzotriazole monohydrate (HOBt, 3.3 mmol, 1.1 equiv.) and, after stirring for 5 min, N-(3-dimethylaminopropyl)-N′-ethylcarbodiimide hydrochloride (EDC·HCl, 3.3 mmol, 1.1 equiv.). The solution was stirred at 0 °C for 1 h. In parallel, L-phenylalanine-L-phenylalanine methyl ester chloride (1.2 g, 3.3 mmol,1.1 equiv.) was dissolved in DCM and treated with N,N-diisopropylethylamine (DIPEA, 9.0 mmol, 3.0 equiv.) added dropwise at room temperature. After stirring for 1 h, this solution was added to the activated Boc-amino acid mixture, rinsing the flask with a small amount of DCM to ensure full transfer. The combined reaction mixture was stirred overnight at room temperature. Upon completion (monitored by TLC, petroleum ether/ethyl acetate 1:2), the reaction mixture was washed sequentially with 0.1 M HCl (3×), then with brine. The organic phase was dried over anhydrous $MgSO_4$, filtered, and concentrated under reduced pressure. The crude product was purified by flash chromatography on silica gel using a gradient elution of petroleum ether/ethyl acetate from 80:20 to 50:50. The pure product was obtained as a white foam (0.74 g, 1.42 mmol, yield: 43%) which was characterized by

NMR. N-Boc-$^{LLL}$PFF-$OCH_3$: $^1H$ NMR (400 MHz, $CDCl_3$, 303 K) δ 7.46 – 6.90 (m, 10H), 4.87 – 4.51 (m,2H), 4.27 – 4.09 (m, 1H), 3.66 (s, 3H), 3.38 – 2.96 (m, 6H), 2.07 (d, J = 15.9 Hz, 1H), 1.76 (m, 3H), 1.43 (s, 9H).

*L-proline-L-phenylalanine-L-phenylalanine methyl ester*: The intermediate compound N-Boc-L-proline-L-phenylalanine-L-phenylalanine methyl ester (0.74 g) was dissolved in 4 mL of 4 M hydrogen chloride in dioxane and stirred at room temperature for 3 hours to allow deprotection. The solvent was then removed under reduced pressure, affording an oily residue. Cold diethyl ether was added to the residue, resulting in the formation of a white precipitate. The supernatant was decanted, and the addition of diethyl ether followed by evaporation was repeated twice to ensure complete removal of residual reagents and solvents. The resulting white solid (0.56 g, yield 99%) was collected and characterized by NMR and mass spectrometry. $^{LLL}$PFF-OMe: $^1H$ NMR (400 MHz, DMSO, 303 K) δ 9.57 (s, 1H), 8.75 (m, 1H), 8.40 (s, 1H), 7.42 – 7.02 (m, 10H), 4.68 – 4.41 (m, 3H), 4.05 (s, 1H), 3.58 (d, *J* = 8.8 Hz, 1H), 3.02 (m, 5H), 2.84 – 2.60 (m, 1H), 2.24 (d, *J* = 6.5 Hz, 1H), 1.99 – 1.57 (m, 4H). HRMS (ESI-Q-Orbitrap): m/z calcd for $C_{24}H_{29}N_3O_4$ $[M+H]^+$ m/z = 424.2231; found m/z = 424.2224.

*N-Boc-D-phenylalanine-D-phenylalanine methyl ester:* N-Boc-D-phenylalanine (1.0 g, 4 mmol, 1.0 equiv.) was dissolved or suspended in dichloromethane (DCM) and cooled to 0 °C. To the resulting mixture were added 1-hydroxybenzotriazole monohydrate (HOBt, 4.4 mmol, 1.1 equiv.) and, after stirring for 5 min, N-(3-dimethylaminopropyl)-N′-ethylcarbodiimide hydrochloride (EDC·HCl, 4.4 mmol, 1.1 equiv.). The solution was stirred at 0 °C for 1 h. In parallel, D-phenylalanine methyl ester hydrochloride (0.93 g, 4.4 mmol, 1.1 equiv.) was dissolved in DCM and treated with N,N-diisopropylethylamine (DIPEA, 12 mmol, 3.0 equiv.) added dropwise at room temperature. After stirring for 1 h, this solution was added to the activated Boc-amino acid mixture, rinsing the flask with a small amount of DCM to ensure full transfer. The combined reaction mixture was stirred overnight at room temperature. Upon completion (monitored by TLC, petroleum ether/ethyl acetate 1:2), the reaction mixture was washed sequentially with 0.1 M HCl (3×), then with brine. The organic phase was dried over anhydrous $MgSO_4$, filtered, and concentrated under reduced pressure. The crude product was purified by flash chromatography on silica gel using a gradient elution of petroleum ether/ethyl acetate from 80:20 to 50:50. The pure product was obtained as a white foam (1.35 g, 3.31 mmol, yield: 83%) which was characterized by NMR. N-Boc-$^{DD}$FF-$OCH_3$:$^1H$ NMR (400 MHz, $CDCl_3$, 303 K) δ 7.64 – 6.86 (m, 10H), 4.86 (dd, *J* = 63.6, 6.6 Hz, 1H), 4.34 (d, *J* = 6.6 Hz, 1H), 3.67 (s, 3H), 3.21 – 2.73 (m, 4H), 1.40 (s, 9H).

*D-phenylalanine-D-phenylalanine methyl ester*: The intermediate compound N-Boc-L-phenylalanine-L-phenylalanine methyl ester (1.35 g) was dissolved in 6 mL of 4 M hydrogen chloride in dioxane and stirred at room temperature for 3 hours to allow deprotection. The solvent was then removed under reduced pressure, affording an oily residue. Cold diethyl ether was added to the residue, resulting in the formation of a white precipitate. The supernatant was decanted, and the addition of diethyl ether followed by evaporation was repeated twice to ensure complete removal of residual reagents and solvents. The resulting white solid (0.96 g, yield: 98%) was collected and characterized by NMR and mass spectrometry. $^{DD}$FF-OMe : $^{1}$H NMR (400 MHz, $D_2O$, 303 K) δ 7.27 – 7.11 (m, 10H), 4.60 (dd, *J* = 7.3. 6.5 Hz, 1H), 4.09 (t, *J* = 7.1 Hz, 1H), 3.58 (s, 3H), 3.10-2.82 (m, 4H). HRMS (ESI-Q-Orbitrap): m/z calcd for $C_{19}H_{22}N_2O_3$ [M+H]+ m/z = 327.1703, found m/z = 327.1699.

*N-Boc-D-proline-D-phenylalanine-D-phenylalanine methyl ester:* N-Boc-D-proline (0.65 g, 3.0 mmol, 1.0 equiv.) was dissolved or suspended in dry dichloromethane (DCM) and cooled to 0 °C. To the resulting mixture were added 1-hydroxybenzotriazole monohydrate (HOBt, 3.3 mmol, 1.1 equiv.) and, after stirring for 5 min, N-(3-dimethylaminopropyl)-N′-ethylcarbodiimide hydrochloride (EDC·HCl, 3.3 mmol, 1.1 equiv.). The solution was stirred at 0 °C for 1 h. In parallel, D-phenylalanine-D-phenylalanine methyl ester chloride (1.2 g, 3.3 mmol,1.1 equiv.) was dissolved in DCM and treated with N,N-diisopropylethylamine (DIPEA, 9.0 mmol, 3.0 equiv.) added dropwise at room temperature. After stirring for 1 h, this solution was added to the activated Boc-amino acid mixture, rinsing the flask with a small amount of DCM to ensure full transfer. The combined reaction mixture was stirred overnight at room temperature. Upon completion (monitored by TLC, petroleum ether/ethyl acetate 1:2), the reaction mixture was washed sequentially with 0.1 M HCl (3×), then with brine. The organic phase was dried over anhydrous $MgSO_4$, filtered, and concentrated under reduced pressure. The crude product was purified by flash chromatography on silica gel using a gradient elution of petroleum ether/ethyl acetate from 80:20 to 50:50. The pure product was obtained as a white foam (0.78 g, 1.49 mmol, yield: 45%) which was characterized by NMR. Boc-$^{DDD}$PFF-OMe $^{1}$H NMR (400 MHz, $CDCl_3$, 303 K) δ 7.41 – 6.91 (m, 10H), 4.81 – 4.56 (m, 2H), 4.33 – 4.02 (m, 1H), 3.67 (s, 3H), 3.32 – 2.84 (m, 6H), 2.19 – 1.77 (m, 4H), 1.43 (s, 9H).

*D-proline-D-phenylalanine-D-phenylalanine methyl ester*: The intermediate compound N-Boc-L-proline-L-phenylalanine-L-phenylalanine methyl ester (0.78 g) was dissolved in 4 mL of 4 M hydrogen chloride in dioxane and stirred at room temperature for 3 hours to allow deprotection. The solvent was then removed under reduced pressure, affording an oily residue. Cold diethyl ether was added to the residue, resulting in the formation of a white precipitate. The supernatant was decanted, and the addition of diethyl ether followed by evaporation was repeated twice to ensure complete removal of residual reagents and solvents. The resulting white solid (0.53 g, yield 99%) was collected and characterized by NMR and mass spectrometry. $^{DDD}$PFF-OMe: 1H NMR (400 MHz, DMSO, 303 K) δ 9.57 (s, 1H), 8.76 (dd, *J* = 27.3, 7.8 Hz, 1H), 8.40 (s, 1H), 7.30 – 6.98 (m, 10H), 4.71 – 4.26 (m,

2H), 4.06 (s, 1H), 3.58 (d, $J$ = 8.8 Hz, 3H), 3.10 – 2.72 (m, 6H), 2.25 (d, $J$ = 6.5 Hz, 1H), 1.78 (ddd, $J$ = 33.8, 16.3, 6.6 Hz, 4H). HRMS (ESI-Q-Orbitrap): m/z calcd for $C_{24}H_{29}N_3O_4$ $[M+H]^+$ m/z = 424.2231; found m/z = 424.2226.

**2.3 NMR Characterization**

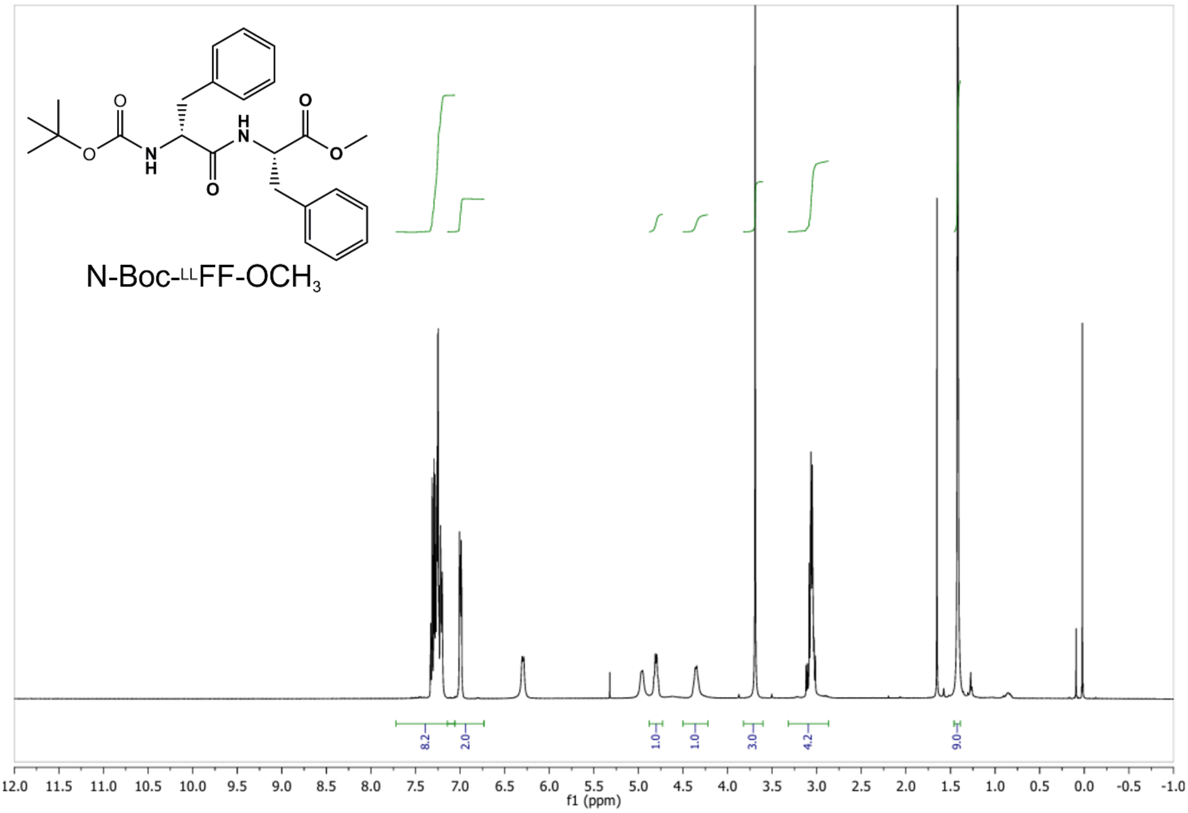

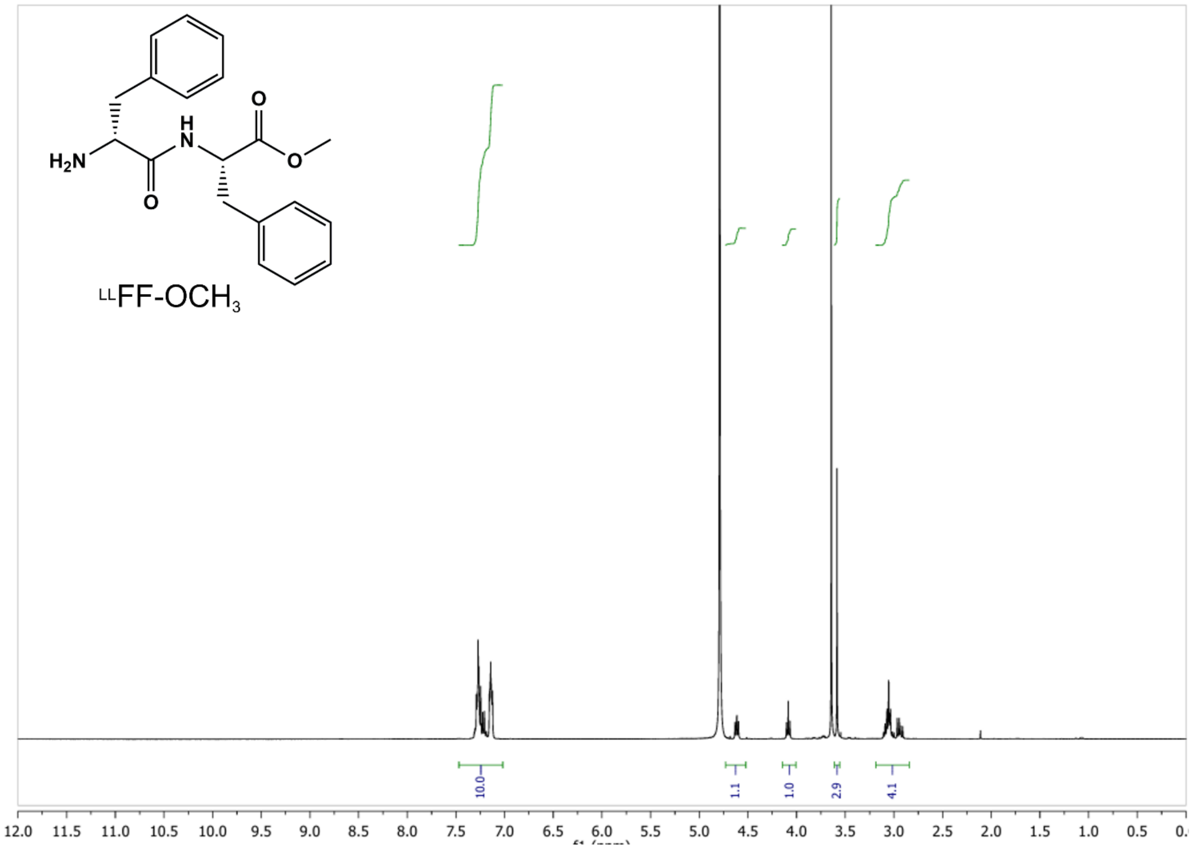

H2N
H
N
O
O
O
LLFF-OCH3
10.0
1.1
1.0
2.9
4.1
12.0 11.5 11.0 10.5 10.0 9.5 9.0 8.5 8.0 7.5 7.0 6.5 6.0 5.5 5.0 4.5 4.0 3.5 3.0 2.5 2.0 1.5 1.0 0.5


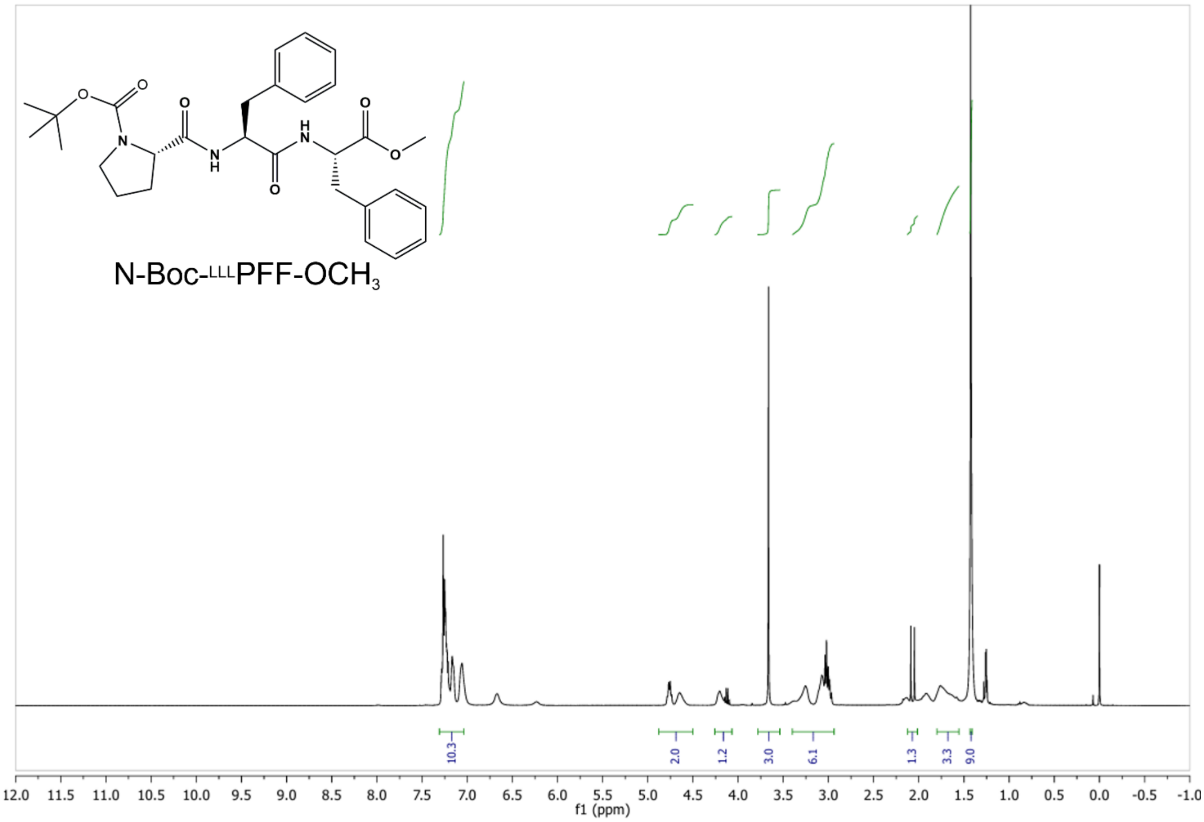

O
O
N
O
N
H
O
H
N
O
O
N-Boc-LLLPFF-OCH3
10.3
2.0
1.2
3.0
6.1
1.3
3.3
9.0
12.0 11.5 11.0 10.5 10.0 9.5 9.0 8.5 8.0 7.5 7.0 6.5 6.0 5.5 5.0 4.5 4.0 3.5 3.0 2.5 2.0 1.5 1.0 0.5 0.0 -0.5 -1.0
f1 (ppm)

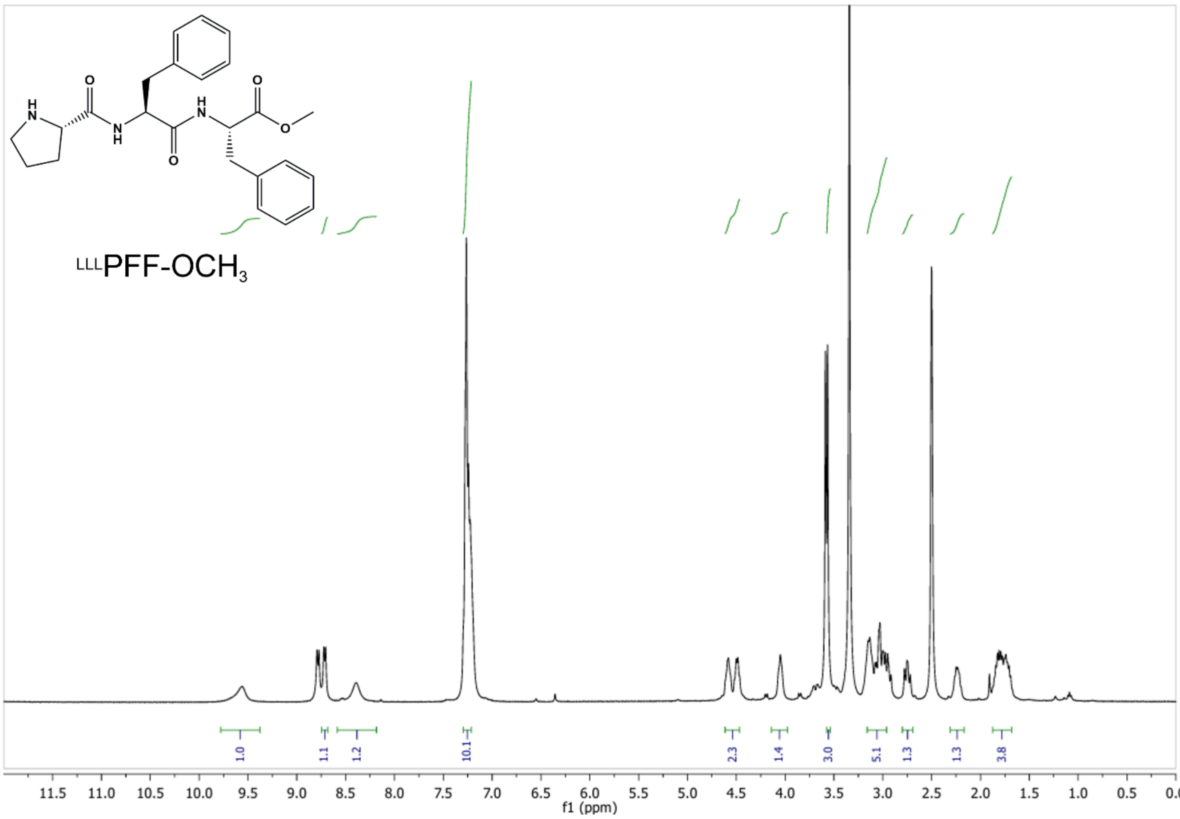
LLLPFF-OCH3
1.0
1.1
1.2
10.1
2.3
1.4
3.0
5.1
1.3
1.3
3.8
11.5 11.0 10.5 10.0 9.5 9.0 8.5 8.0 7.5 7.0 6.5 6.0 5.5 5.0 4.5 4.0 3.5 3.0 2.5 2.0 1.5 1.0 0.5 0.0
f1 (ppm)

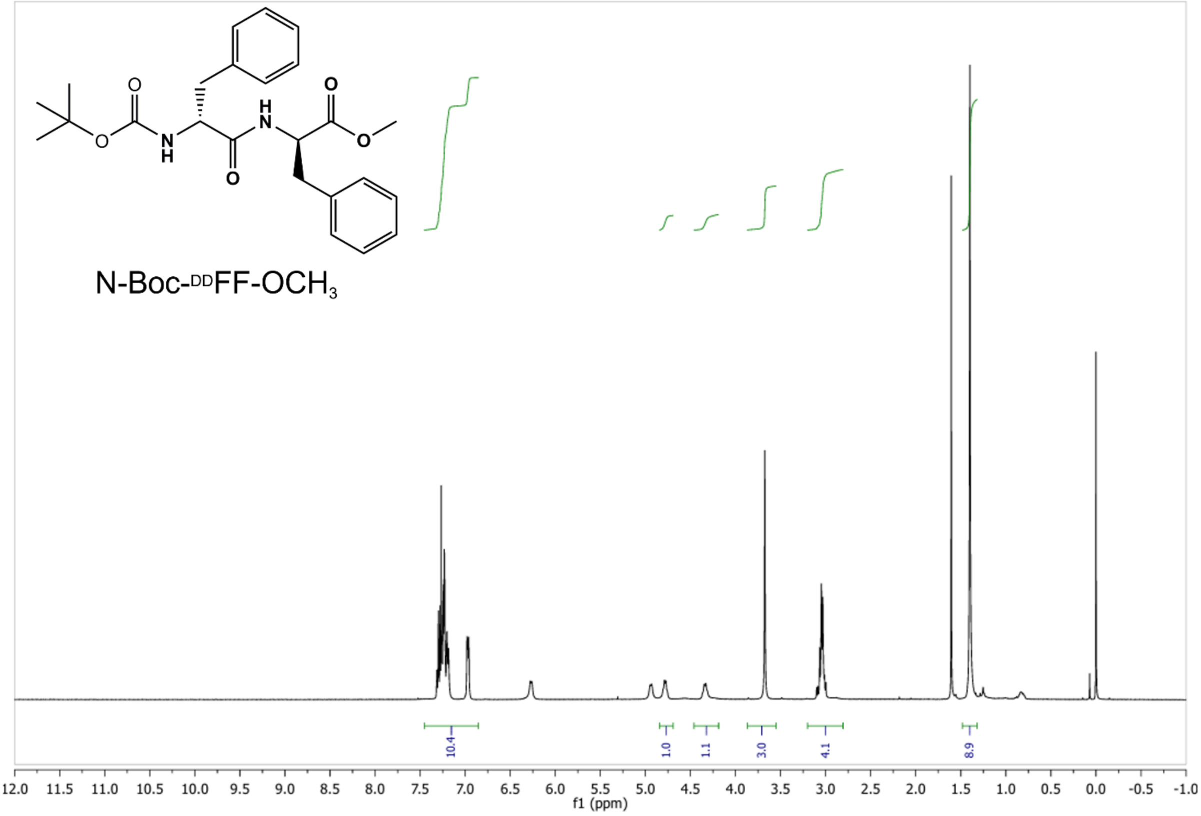
N-Boc-DDFF-OCH3
10.4
1.0
1.1
3.0
4.1
8.9
12.0 11.5 11.0 10.5 10.0 9.5 9.0 8.5 8.0 7.5 7.0 6.5 6.0 5.5 5.0 4.5 4.0 3.5 3.0 2.5 2.0 1.5 1.0 0.5 0.0 -0.5 -1.0
f1 (ppm)

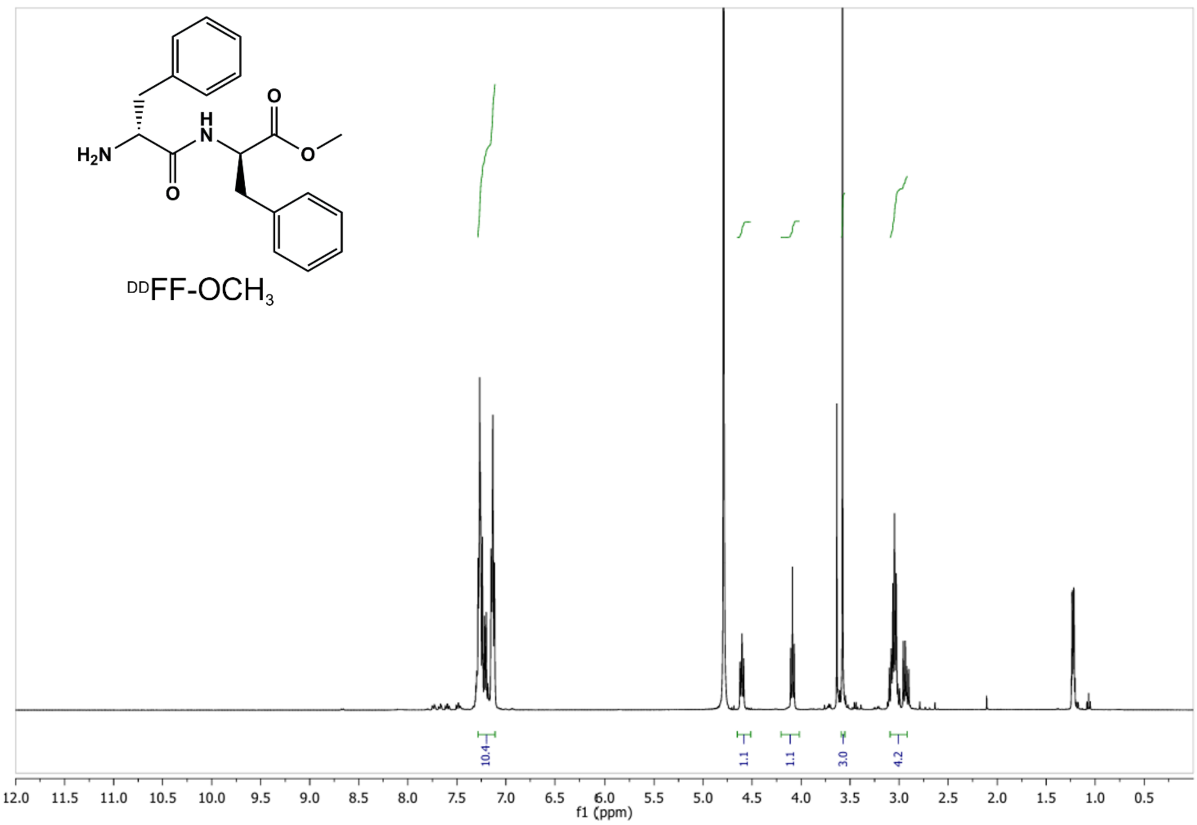

DDFF-OCH3
10.4
1.1
1.1
3.0
4.2
f1 (ppm)


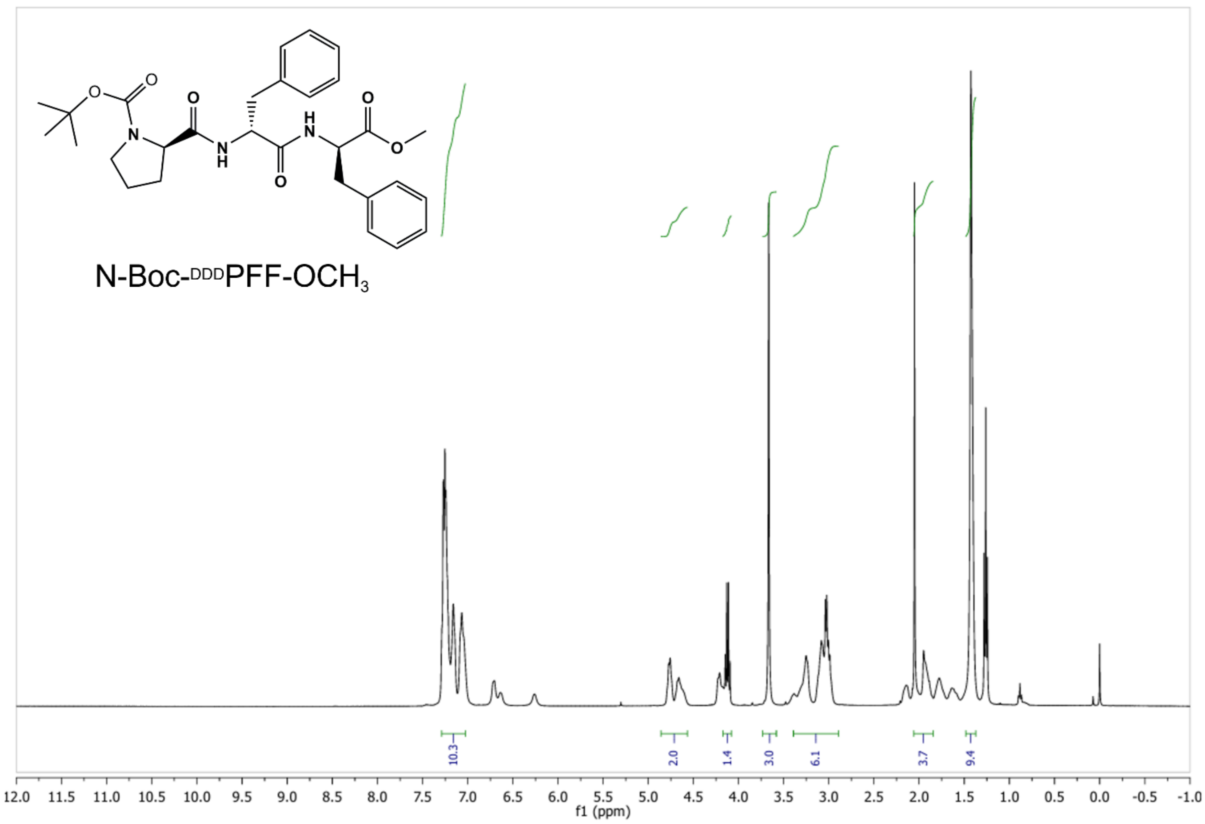

N-Boc-DDDPFF-OCH3
10.3
2.0
1.4
3.0
6.1
3.7
9.4
f1 (ppm)

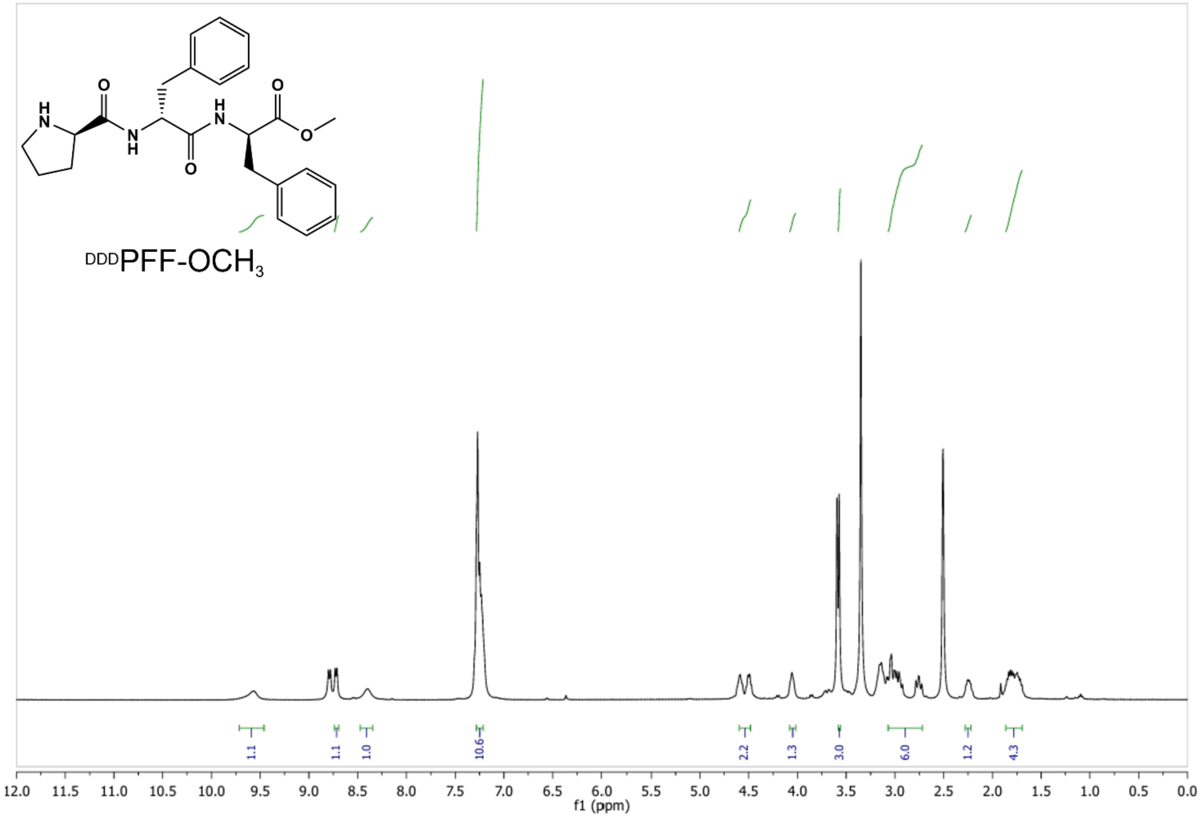


#### 2. Peptide Coacervates: Preparation and Functional Properties

##### 3.1 Coacervate Preparation

PFF-$OCH_3$ and FF-$OCH_3$ were dissolved at the desired concentration, e.g. 10 mg·$mL^{-1}$, in 50 mM phosphate buffer (pH 6). The pH was adjusted to the desired value by the addition of 3 M NaOH solution. All vials were wetted with 1% Pluronic® F-108 and then rinsed with the appropriate phosphate buffer before use to prevent peptide adhesion.

##### 3.2 Turbidity Measurements

Absorbance at 600 nm was measured using a Microvolume Spectrophotometer (VWR). Turbidity was used as an indicator of phase separation, and droplet formation was confirmed by confocal microscopy. Absorbance at 600 nm was recorded for all measurements. Unless otherwise stated, measurements were carried out at room temperature. After sample loading and brief equilibration, turbidity values were recorded. 50 mM phosphate buffer was used as blank for baseline correction.

##### 3.3 Guest Molecule Partitioning

"1 µL of a 1 mg·mL$^{-1}$ aqueous Rhodamine B solution was added to 20 µL of a turbid 10 mg·mL$^{-1}$ peptide solution prepared in 50 mM phosphate buffer at pH 8.5. The mixture was then transferred into a µ-Slide 8 Well (Ibidi) and mounted directly onto the microscope stage. Droplet imaging was carried out by confocal microscopy using a Leica SP8 Stellaris Falcon confocal microscope equipped with HyD photon counting detectors and a white light laser (WLL) excitation source, tunable from 405 to 850 nm. Image analysis was performed using LAS X software (Leica Microsystems).

##### 3.4 D-SPR analysis

D-SPR experiments were performed on a Bionavis SPR Navi 210A multiparametric instrument. This system is equipped with 70 cm long injection tubes and features two parallel channels, along with two distinct laser wavelengths (670 nm and 750 nm). For all analyses, only the 670 nm laser was used. The experiments were run on an SPR102-AU gold sensor chip 20 mm long, 12 mm wide, and 0.55 mm thick; the area in contact with the flow is 12 mm$^2$. Before starting the diffusion experiments, the instrument was set to record a response point every 1.8 s, and 50 mM phosphate buffer injections were performed to stabilize the baseline. Each peptide sample solution was injected at a flow rate of 10 µl·min$^{-1}$ four times, with the first injection serving to condition the microfluidic system and confirm that there is no interaction between the analyte and the surface. Coacervate characterization analyses were performed at 26°C. All the data collected was processed with the Origin 2018 software®. The determination of the diffusion coefficient was conducted using a dedicated Python-based script as previously reported.[3] The code is available upon request from the authors.

#### 3. Characterization of Coacervates

##### 4.1 Fluorescence Confocal Microscopy

Fluorescence imaging was performed on a Leica SP8 Stellaris Falcon confocal microscope equipped with HyD photon counting detectors and a white light laser (WLL) excitation source, tunable from 405 to 850 nm. A HC PL APO CS2 63×/1.40 NA oil immersion objective was used. The electronic zoom was set to 3.0×, resulting in a pixel size of 0.03 µm, and images were acquired at a resolution of 2048 × 2048 pixels, corresponding to a field of view of 61.42 × 61.42 µm. The pinhole diameter was set to 95.5 µm (1 Airy unit, calculated for 580 nm emission). Excitation was performed at 543 nm, and emission was detected between 605 and 637 nm using the HyD S 2 detector. Scanning was performed in xyt mode at 700 Hz in unidirectional mode, with a pixel dwell time of

0.375 µs. Samples were transferred to a µ-Slide 8 Well (Ibidi) and mounted directly onto the microscope stage at room temperature.

##### 4.2 FRAP Measurements

For FRAP experiments, 20 µL of the peptide coacervate solution prepared as described above was mixed with 1 µL of a Rhodamine B solution (1 mg·mL$^{-1}$ in water). The sample was transferred into a µ-Slide 8 Well (Ibidi) and mounted on the microscope stage. After thermal equilibration, the FRAP Wizard mode was activated using the same 63× oil immersion objective. Images were acquired using HyD fluorescence detection only, at a resolution of 256 × 256 pixels, with an electronic zoom of 6×, resulting in a final pixel size of 0.12 µm and a field of view of approximately 30.72 × 30.72 µm. The 543 nm laser line was used for both imaging and photobleaching. The FRAP acquisition protocol consisted of a 20 s pre-bleach phase to establish baseline fluorescence, followed by a 2 s bleaching pulse in a defined region of interest, and a 150 s post-bleach phase to monitor fluorescence recovery. The bleaching intensity was adjusted to achieve a fluorescence drop of approximately 50% relative to the initial intensity immediately after bleaching. All acquisition parameters were kept constant throughout the experiments. Image analysis was performed using LAS X software (Leica Microsystems).

##### 4.3 Micro-Raman Spectroscopy

Micro-Raman measurements were carried out using a confocal Raman microscope (HORIBA Jobin Yvon Labram) equipped with a CW laser operating at 532 nm with an output power of 50 mW. The excitation light was focused onto the sample using a dry objective lens (Olympus LMPlanFl N 50x/0.5 NA) mounted on an upright microscope, and Raman-scattered light was then collected in backscattering geometry through the same objective. Diffused light was then dispersed by a 1800 gr/mm diffraction grating and detected using a front-illuminated, thermoelectrically cooled CCD camera (Horiba SYNAPSE, 1024 × 256 pixels). The system provides an in-plane spatial resolution of approximately 1.5 µm and a spectral resolution of 1.34 cm$^{-1}$. Prior to measurements, the spectrometer was calibrated using the first-order Raman peak of a Silicon standard, aligning it at 521 cm$^{-1}$ to ensure accurate spectral positioning. Only experiments aimed at analyzing the Amide I signals were performed under identical conditions, except that $D_2O$ was used instead of $H_2O$. Additionally, spectra were collected on the solid racemic precipitate.

Estimation of $^{LLL}$PFF-$OCH_3$ concentration in the coacervates

Raman intensity ratios (R) between the $^{LLL}$PFF-$OCH_3$-specific band at ~ 1003 $cm^{-1}$ (F5)[4] and the solvent-specific band at ~ 3400 $cm^{-1}$ (water O-H asymmetric stretching) provide a quantity proportional to the pro-F-F concentration (C) in a sampled region. Based on the intensity ratio observed for the transparent 10 mg·$mL^{-1}$ solution of $^{LLL}$PFF-$OCH_3$ at pH 6, we estimated the concentration of $^{LLL}$PFF-$OCH_3$ in the $^{LLL}$PFF-$OCH_3$ droplets at pH 8.5, considering the following proportional relationship:

$$C_B : R_B = C_A : R_A$$

where 'A' and 'B' refer pH 6 and coacervates at pH 8.5 respectively.

The aforementioned ratios were measured inside multiple droplets, providing a statistically meaningful estimate of the concentration value and its associated standard deviation. The mean value inside the droplet is given by:

$$\overline{C}_{\mathrm{D}} = \frac{C_{\mathrm{A}}}{R_{\mathrm{A}}} \cdot \frac{1}{N} \sum_{droplets}^{N} R_{\mathrm{D}} = 347\ \mathrm{mg \cdot mL^{-1}}$$

whereas the standard deviation was calculated considering error propagation over the intensity ratios:

$$\sigma_{C_D} = \frac{C_A}{\mathrm{R}_A} \cdot \sigma_{R_D} = 28\ \mathrm{mg \cdot mL^{-1}}$$

**4.4 IR Spectroscopy**

Fourier Transform Infrared (FTIR) spectra were acquired using a Bruker VERTEX 70v spectrometer equipped with a Globar MIR source, a DLaTGS detector with CsI window, and a single-reflection diamond crystal ATR accessory (Platinum ATR, Bruker). Spectra were recorded in ATR mode over the spectral range of 4000–400 $cm^{-1}$ with a resolution of 0.5 $cm^{-1}$. For each acquisition, 128 scans were co-added and averaged using OPUS 7.8 software. Background spectra were recorded under the same conditions and subtracted from each sample spectrum. Measurements on solid samples were performed under vacuum to reduce atmospheric interference, while liquid samples were measured under ambient pressure. The analysis was performed on a 10 mg·$mL^{-1}$ solution of $^{LLL}$PFF-$OCH_3$ in $D_2O$ at pH 6 and 8. Additionally, spectra were collected on the solid racemic precipitate.

#### 4. Computational Analysis

##### 5.1 Molecular Dynamic Details

MD simulations were performed with the GROMACS package (version 5.1.2) in the NPT ensemble using a cuboidal simulation box, the velocity rescaling temperature coupling, and the Parrinello–Rahman barostat with 2 ps relaxation times. Periodic boundary conditions were used, and the long-range electrostatic interactions were treated with the particle mesh Ewald method with a real space cutoff of 0.9 nm. The Lennard-Jones potential was truncated at 0.9 nm. The LINCS algorithm was used to constrain bond lengths along with a 2 fs time step. The CHARMM36m force field was employed to model the peptide.

##### 5.2 Computational Analyses

**Hbond Analysis.** Hydrogen bond (H-bond) analysis was performed on the MD trajectories using the gmx hbond tool from the GROMACS package. H-bonds were identified based on a donor–acceptor distance cutoff of 0.35 nm and a hydrogen–donor–acceptor angle cutoff of 30°, using default geometric criteria. The analysis was conducted over the entire production trajectory to evaluate the average number and stability of H-bonds formed between relevant molecular groups.

**End-To-End Distance.** The end-to-end distance of the peptide was calculated as the distance between the carbon atom of the methyl group of the phenylalanine and the nitrogen of the N-terminal proline over the course of the MD trajectory. This analysis was performed using MDAnalysis, and the resulting distance values were monitored to assess the conformational flexibility and overall extension of the peptide during the simulation.

**Ramachandran plot.** Ramachandran plot analysis was performed to evaluate the backbone dihedral angle distributions ($\varphi$ and $\psi$) of the peptide during the simulation. The analysis was carried out using the gmx rama tool in GROMACS over the production trajectory, allowing for the identification of preferred conformational regions and the assessment of structural flexibility and secondary structure tendencies.

**Radial Distribution Functions.** Radial distribution function (RDF) analysis was performed using the gmx rdf tool in GROMACS to investigate the spatial distribution of water around nitrogen atoms on the peptide. The RDF was calculated over the production trajectory to quantify local structural organization and intermolecular interactions. Peaks in the RDF were used to identify preferred interaction distances and coordination environments.

**Solvent Accessible Solvent Area (SASA).** Solvent-accessible surface area (SASA) analysis was performed using the gmx sasa tool in GROMACS to quantify the exposure of the peptide to the solvent. To identify the different aminoacid contributions to the SASA another analysis is performed, considering in a separate way the aromatic rings of the F-F and the pyrrolidine ring of the proline.

**F-F intramolecular and intermolecular distances.** The distance between the para carbon atoms (Cε) of the phenyl rings of phenylalanine residues was calculated to assess π–π interactions. Both intramolecular and intermolecular distances were measured using MDAnalysis. Atom selections were based on the Cε,, and the analysis was performed over the production trajectory to evaluate the entity and geometry of potential stacking interactions.

**Clustering with DBSCAN.** Clustering analysis was performed using the DBSCAN algorithm to identify phenylalanine side-chain stacking arrangements. The analysis focused on the para carbon atoms (Cε) of the phenyl rings. The distance threshold (ε) for DBSCAN was selected based on the first minimum following the first peak in the radial distribution function (RDF) of para carbon–para carbon distances, computed from the all-neutral simulation. A minimum number of points (minPts) equal to 3 was used to detect clusters in which a para carbon atom is surrounded by at least two neighboring para carbon atoms within the RDF-derived cutoff. This approach allowed the identification of both intra- and intermolecular π–π stacking assemblies.

| | Avg. Biggest Cluster Dimension | Avg. Number of Clusters | Avg. Cluster Dimension |
|---|---|---|---|
| Cation | 47 | 41 | 11 |
| 93:7 | 59 | 37 | 13 |
| Neutral | 387 | 13 | 47 |

**Table S1.** Summary of clustering analysis using the DBSCAN algorithm, considering the para carbon atoms (Cε) of the phenylalanine rings as density points. Reported values include the average size of the largest cluster (as a percentage of total peptides in the simulation box), the average number of clusters per frame, and the average cluster size (also as a percentage of total peptides).

#### 5. Aldol Reaction in Coacervates

##### 6.1 General Methods

$^1$H NMR spectra were recorded at 295 K on a Bruker Avance III 400 MHz spectrophotometer. Chemical shifts (δ) are reported in parts per million (ppm) relative to the residual solvent signal ($^1$H NMR: 7.26 ppm for $CDCl_3$). Coupling constants (*J*) are given in hertz (Hz). $^1$H NMR yields were determined by analysing the crude reaction mixtures using triphenylmethane as internal standard. Abbreviations used in signal descriptions are as follows: s, singlet; d, doublet; t, triplet; q, quartet; p, pentet; br s, broad singlet. The diastereomeric ratio was determined by $^1$H-NMR of the crude product. The enantiomeric excess (*ee*) of the aldol product was determined by chiral HPLC using a hexane/2-propanol mixture as the mobile phase. Analyses were performed on an Agilent 1220 Infinity II liquid chromatograph equipped with a Phenomenex Lux Cellulose-1 column (100 × 4.6 mm, 3 µm particle size). HPLC traces were compared to a racemic sample prepared using a 10 M sodium hydroxide solution, in water for 16 h [*Organic & Biomolecular Chemistry (2013) 11 2951–2960*]. The absolute configurations of the optically active compounds were determined by comparing the measured optical rotations with literature values [*Journal of the American Chemical Society (2003) 125(9) 2475–2479*]. Measurements were performed using a ZUZI 412 Digital Polarimeter equipped with a 100 mm tube. Percoated ALUGRAM Xtra SIL G/UV 254 silica gel plates were used for thin layer analytical chromography (TLC) and visualized with short wave UV light or by *p*-anisaldehyde stain. Column chromatography was carried out employing EMD (Merck) Silica Gel 60 (35-70 µm). All starting materials were purchased from Merck and used without any further purification. HPLC solvents and analytical grade solvents were purchased from Merck and used as received (no extra drying, distillation or special handling practices were employed).

#### 6.2 Experimental procedures

*Reaction scheme for the peptide-catalysed aldol reaction*

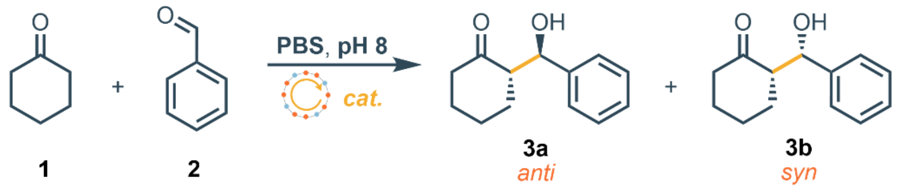


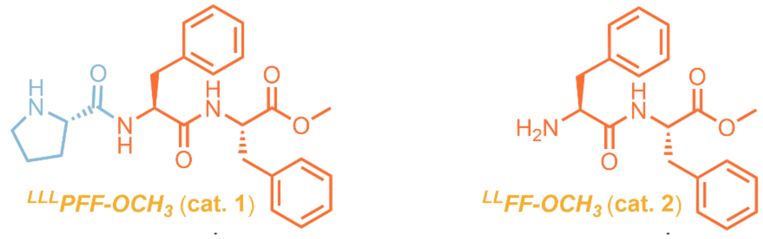


##### Synthesis of the racemic sample

A stirred solution of water (20 mL) was charged with 10 M NaOH (1 mL). Cyclohexanone (0.28 mL, 2.7 mmol, 5 equiv.) was added and the solution was stirred for 1 min. To the mixture was added benzaldehyde (0.05 mL, 0.54 mmol, 1 equiv.) and the solution was stirred overnight at room temperature. The mixture was extracted into $CHCl_3$ (3 × 20 mL), and the combined organic phase was dried with $Na_2SO_4$. Purification over silica gel eluting in Petroleum/EtOAc 9:1 afforded the two diastereomers **3a** and **3b**.

##### General Procedure for the peptide-organocatalysed aldol reaction

Benzaldehyde (11.6 mg, 0.1 mmol, 1 equiv.) and cyclohexanone (21.4 mg, 0.2 mmol, 2 equiv.) were added to 1 mL of a PBS aqueous solution containing the peptide catalyst (10 mg·$mL^{-1}$, 0.02 mmol, 0.2 equiv). The mixture was stirred for 20 hours at 4 °C. THF (2 mL) was added to the reaction mixture to disperse the droplets. The crude mixture was then filtered through a short silica plug (DCM/$Et_2O$ 1:1) and the resulting solution was concentrated *in vacuo*. Purification over silica gel eluting in Petroleum/EtOAc 9:1 afforded the pure product **3a**.

##### Characterisation data

*(S)-2-((R)-hydroxy(phenyl)methyl)cyclohexan-1-one* (**3a**) **$^1$H NMR** (400 MHz, $CDCl_3$) δ 7.38 – 7.27 (m, 5H), 4.79 (d, *J* = 8.8 Hz, 1H), 3.96 (br s, 1H), 2.67 – 2.57 (m, 1H), 2.52 – 2.45 (m, 1H), 2.41 – 2.31 (m, 1H), 2.13 – 2.04 (m, 1H), 1.83 – 1.74 (m, 1H), 1.73 – 1.48 (m, 4H), 1.36 – 1.24 (m, 1H). HPLC analysis: Lux 3 µm-Cellulose 1, Hexane/i-Propanol 90:10, flow 0.5 mL/min, 210 nm, 25 °C): $t_{minor}$ = 10.3 min, $t_{major}$ = 8.3 min. $[\alpha]_D^{25}$ = + 23.3

(c=0.0006 g/mL, $CHCl_3$). Analytical data in accordance with literature report [*European Journal of Organic Chemistry (2014) 7 624-630*].

*(S)-2-((S)-hydroxy(phenyl)methyl)cyclohexan-1-one* (**3b**) **$^1$H NMR** (400 MHz, $CDCl_3$): δ = 7.42 – 7.22 (m, 5H), 5.39 (d, $J$ = 2.5 Hz, 1H), 3.03 (br s, 1H), 2.63 - 2.57 (m, 1H), 2.50 – 2.27 (m, 2H), 2.12 – 2.04 (m, 1H), 2.13 – 2.05 (m, 1H), 1.87 – 1.81 (m, 1H), 1.77 - 1.61 (m, 3H), 1.57 – 1.44 (m, 1H). Analytical data in accordance with literature report [*Organic & Biomolecular Chemistry (2011) 9 1784-1790*]

**NMR spectra**

*$^1$H NMR spectrum (in $CDCl_3$) of (S)-2-((R)-hydroxy(phenyl)methyl)cyclohexan-1-one (**3a**)*

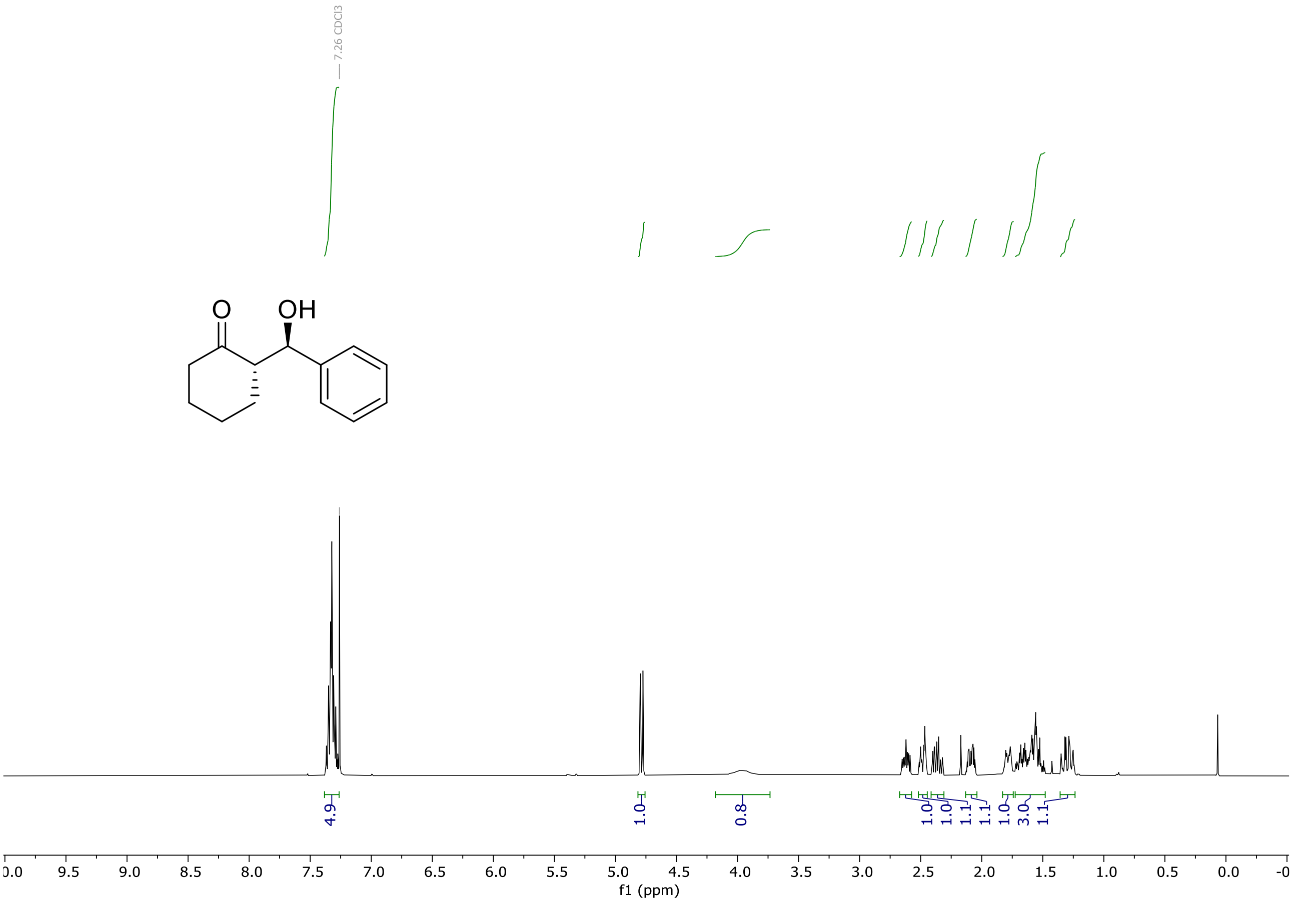

*[1]H NMR spectrum (in $CDCl_3$) of (S)-2-((S)-hydroxy(phenyl)methyl)cyclohexan-1-one (**3b**)*

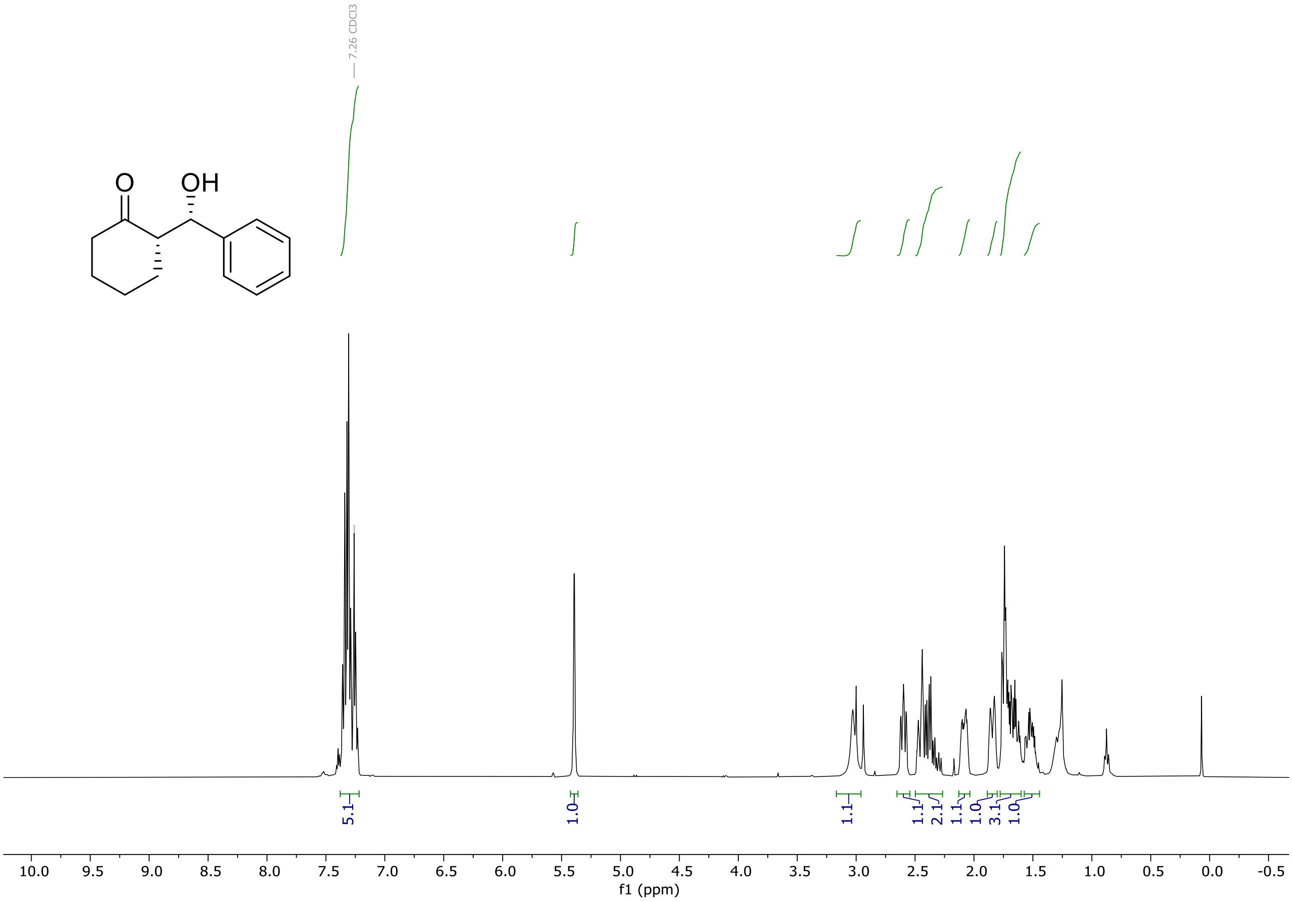


**HPLC Traces**

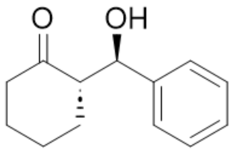


Lux 3 µm-Cellulose 1, Hexane/i-Propanol 90:10, flow 0.5 mL/min, λ= 210 nm

**Racemate**

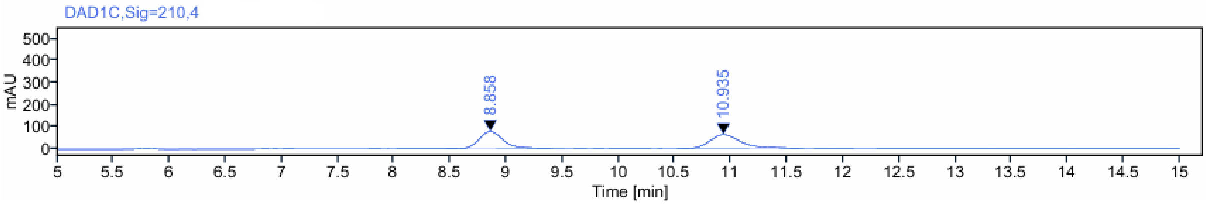

| Signal: | DAD1C,Sig=210,4 | | | | |
|---|---|---|---|---|---|
| RT [min] | Type | Width [min] | Area | Height | Area% |
| 8.858 | MM m | 0.23 | 1158.82 | 77.66 | 49.32 |
| 10.935 | MM m | 0.29 | 1190.65 | 62.01 | 50.68 |
| | | Sum | 2349.48 | | |

**Product (3a) obtained employing cat. 1**

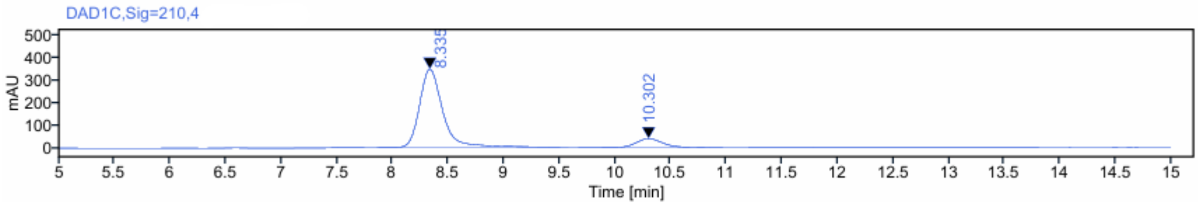


| Signal: | DAD1C,Sig=210,4 | | | | |
|---|---|---|---|---|---|
| RT [min] | Type | Width [min] | Area | Height | Area% |
| 8.335 | MM m | 0.22 | 4989.28 | 344.22 | 89.06 |
| 10.302 | MM m | 0.25 | 613.07 | 38.22 | 10.94 |
| | | Sum | 5602.35 | | |

**Product (3a) obtained employing cat. 2**

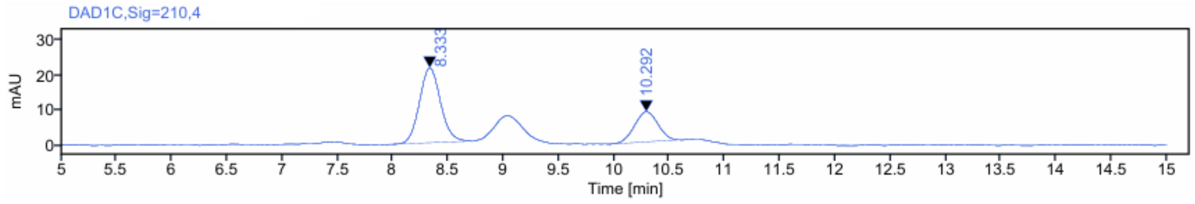


| Signal: | DAD1C,Sig=210,4 | | | | |
|---|---|---|---|---|---|
| RT [min] | Type | Width [min] | Area | Height | Area% |
| 8.333 | MM m | 0.21 | 280.24 | 21.17 | 69.11 |
| 10.292 | BM m | 0.24 | 125.25 | 8.48 | 30.89 |
| | | Sum | 405.50 | | |
